# Benchmarking ideal sample thickness in cryo-EM using MicroED

**DOI:** 10.1101/2021.07.02.450941

**Authors:** Michael W. Martynowycz, Max T.B. Clabbers, Johan Unge, Johan Hattne, Tamir Gonen

## Abstract

The relationship between sample thickness and quality of data obtained by microcrystal electron diffraction (MicroED) is investigated. Several EM grids containing proteinase K microcrystals of similar sizes from the same crystallization batch were prepared. Each grid was transferred into a focused ion-beam scanning electron microscope (FIB/SEM) where the crystals were then systematically thinned into lamellae between 95 nm and 1650 nm thick. MicroED data were collected at either 120, 200, or 300 kV accelerating voltages. Lamellae thicknesses were converted to multiples of the calculated inelastic mean free path (MFP) of electrons at each accelerating voltage to allow the results to be compared on a common scale. The quality of the data and subsequently determined structures were assessed using standard crystallographic measures. Structures were reliably determined from crystalline lamellae only up to twice the inelastic mean free path. Lower resolution diffraction was observed at three times the mean free path for all three accelerating voltages but the quality was insufficient to yield structures. No diffraction data were observed from lamellae thicker than four times the calculated inelastic mean free path. The quality of the determined structures and crystallographic statistics were similar for all lamellae up to 2x the inelastic mean free path in thickness, but quickly deteriorated at greater thicknesses. This study provides a benchmark with respect to the ideal limit for biological specimen thickness with implications for all cryo-EM methods.

**Significance:** A systematic investigation of the effects of thickness on electron scattering from protein crystals was previously not feasible, because there was no accurate method to control sample thickness. Here, the recently developed methods for preparing protein crystals into lamellae of precise thickness by ion-beam milling are used to investigate the effects of increasing sample thickness on MicroED data quality. These experiments were conducted using the three most common accelerating voltages in cryo-EM. Data across these accelerating voltages and thicknesses were compared on a common scale using their calculated inelastic mean free path lengths. It is found that structures may accurately be determined from crystals up to twice the inelastic mean free path length in thickness, regardless of the acceleration voltage.

## Main

High energy electrons interact strongly with matter (1–3). This property has enabled a revolution in structural biology by electron cryo-microscopy (cryo-EM) techniques (4). However, this strong interaction also implies a higher probability of an electron scattering multiple times and/or losing energy within the specimen (5). The probability of scattering relates to a physical property known as the mean free path. This is the average distance travelled through a sample by a moving particle before an interaction takes place. The inelastic mean free path (MFP) refers to the typical distance a high-energy electron travels through a specimen before losing energy, or inelastically scattering. In cryo-EM, the MFP is often used to compare samples of different thicknesses across different accelerating voltages (6, 7). The MFP in cryo-EM may be roughly calculated for a given sample, and has been investigated experimentally in vitreous ice, since this is the most probable environment in these experiments, though similar values have recently been demonstrated in liquid water (7, 8). The calculated MFP for a typical protein crystal at accelerating voltages of 120, 200 or 300kV would correspond to roughly to 214, 272 and 317 nm, respectively (Materials and Methods).

Early cryo-EM investigations of electron diffraction from frozen-hydrated protein samples reported measurable differences between the intensities of Friedel mates from two-dimensional crystals of bacteriorhodopsin (bR) (9). These differences were suggested to arise from dynamically scattered electrons. Dynamically scattered electrons could introduce significant errors, breaking the relationship between the recorded intensity and the underlying structure factor amplitude. Computational simulations suggested that two-dimensional crystals of bR thicker than 20 nm at 100 kV, and three-dimensional crystals of lysozyme thicker than 100 nm at 200 kV would result in highly inaccurate intensities due to dynamical scattering (10, 11). Those results are at odds with earlier reports that diffraction intensities from three-dimensional catalase crystals at 200 kV scatter kinematically at thicknesses up to at least 150 nm (12). Indeed, investigations reported structures of catalase from crystals of variable crystal thicknesses without the need of any dynamical corrections (13, 14). Many macromolecular structures have since been reported from crystals that are thicker than 100 nm using the cryo-EM method microcrystal electron diffraction (MicroED) (14–20).

Until recently, a systematic investigation of how crystal thickness effects data quality was not feasible. Now, FIB milling allows the thickness of a protein crystal to be precisely controlled (21–26). This process is similar to milling cellular and tissue specimens to prepare them for subsequent cryo-tomography investigations (21, 23, 27, 28). Zhou et al. recently used this technique to mill several crystals to different thicknesses and compared single diffraction pattern from each at 200 kV (24). However, they did not systematically correlate the effect of crystal thickness on the ability to determine structures. Only a systematic investigation of integrated intensities and the quality of the determined protein structures from lamellae of variable thicknesses would shed light on the role of sample thickness in cryo-EM and possible errors due to dynamical scattering.

Here, we investigate the impact of sample thickness on the ability to determine structures and the quality of data obtained using MicroED. Microcrystals of proteinase K were milled into lamellae between 95 and 1650 nm thick. MicroED data were collected from each lamella at one of the three most common acceleration voltages (120, 200, and 300 kV) (Figure 1). The data were put on a common scale by relating their thicknesses to the inelastic mean free path at their respective acceleration voltage. These thicknesses roughly correspond to between 0.5× and 5× MFP. We found that MicroED data from crystals as thick as twice the MFP are sufficiently accurate to determine high-resolution protein structures irrespective of the acceleration voltage. Surprisingly, no large difference in data or structure quality was observed from lamellae below 2× MFP. Diffraction was still observed at up to 3× MFP, but the data was not suitable for processing. No diffraction spots were observed for thicknesses beyond 4× MFP. These trends were true for all three acceleration voltages. This study provides initial measurements of crystals of definitive thicknesses at varying accelerating voltages and provides limits on biological specimen thickness with implications for all cryo-EM investigations.

**Figure 1.**
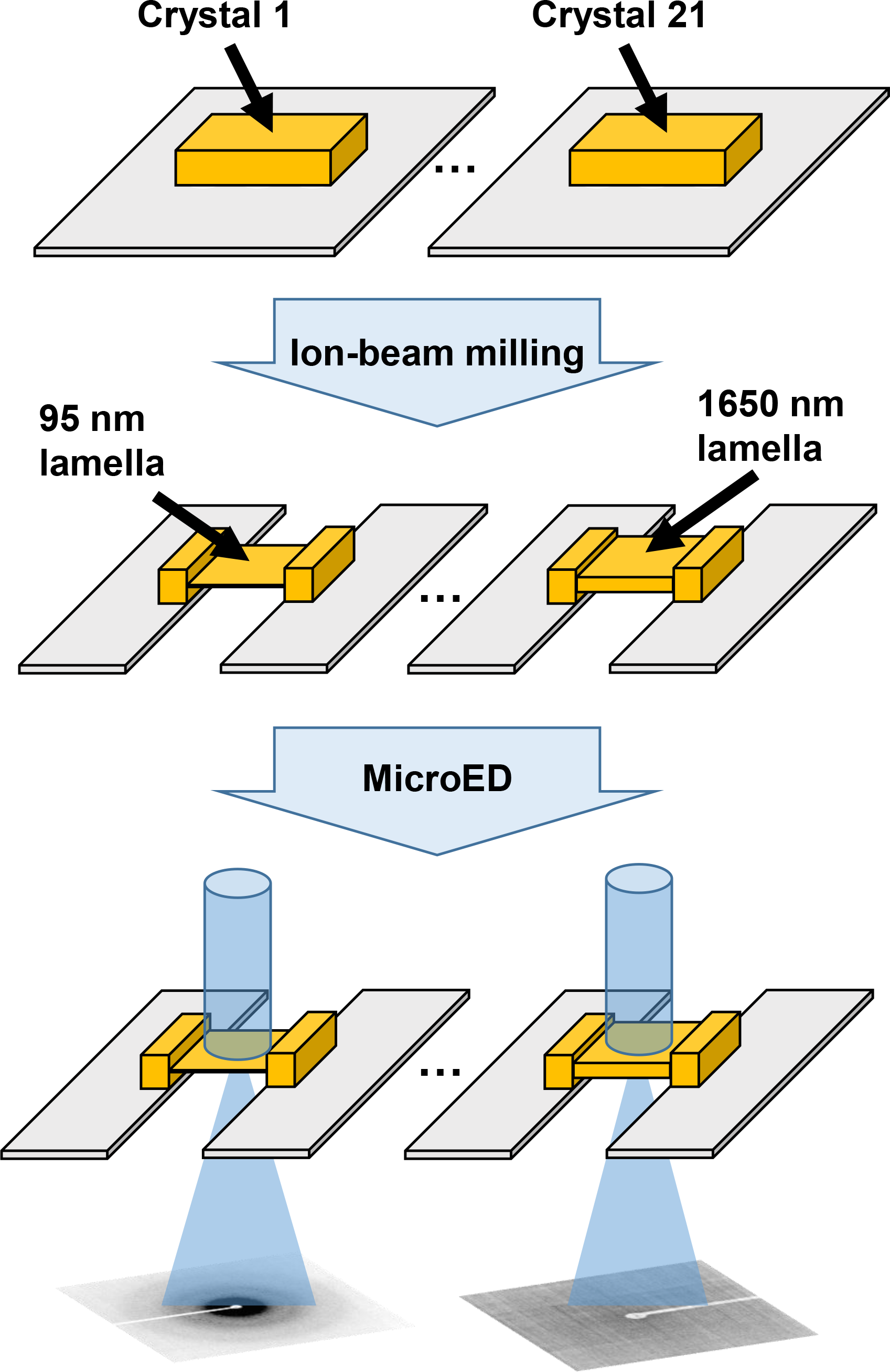
Preparation of protein microcrystals into lamellae of specified thicknesses. Schematic cartoon showing the general process of systematically investigating data quality for variably thick samples. Crystals are identified on EM grids (top), milled to specified thicknesses (middle), and MicroED datasets are collected from each crystal at either 120, 200, or 300 kV accelerating voltages.

## Results

### Preparing grids with protein crystals

Proteinase K crystals were grown in batch as described (29, 30). This condition results in protein microcrystals that measure between 10 and 30 µm across their middle. TEM grids were prepared by back blotting as described (30). The samples were then loaded into a focused ion-beam scanning electron microscope (FIB/SEM).

Vitrified grids were coated with sputtered platinum to protect the crystals from the damaging electron and ion-beams during investigation (29) (Materials and Methods). Each grid was searched using the scanning electron beam for crystals that satisfied the following requirements: each crystal was of relatively similar size, was at least 5 µm away from the nearest grid bar, and at least 3 grid squares away from the edge of the grid. In this way, 5 appropriate crystals were identified on a first batch of grids, 9 on the second, and 7 on the third. Each crystal was inspected in both the SEM and FIB and aligned to eucentric height. Lamellae were milled by rastering the gallium beam across the microcrystals (Materials and Methods). The ion-beam current was lowered as the desired thickness was approached as described (27, 29). The thickness of each crystalline lamella was measured by taking an image in the ion-beam just prior to unloading each grid. Lamellae thicknesses spanned from approximately 95 to 1650 nm (Figure 1, Figure 2) (Materials and Methods).

**Figure 2.**
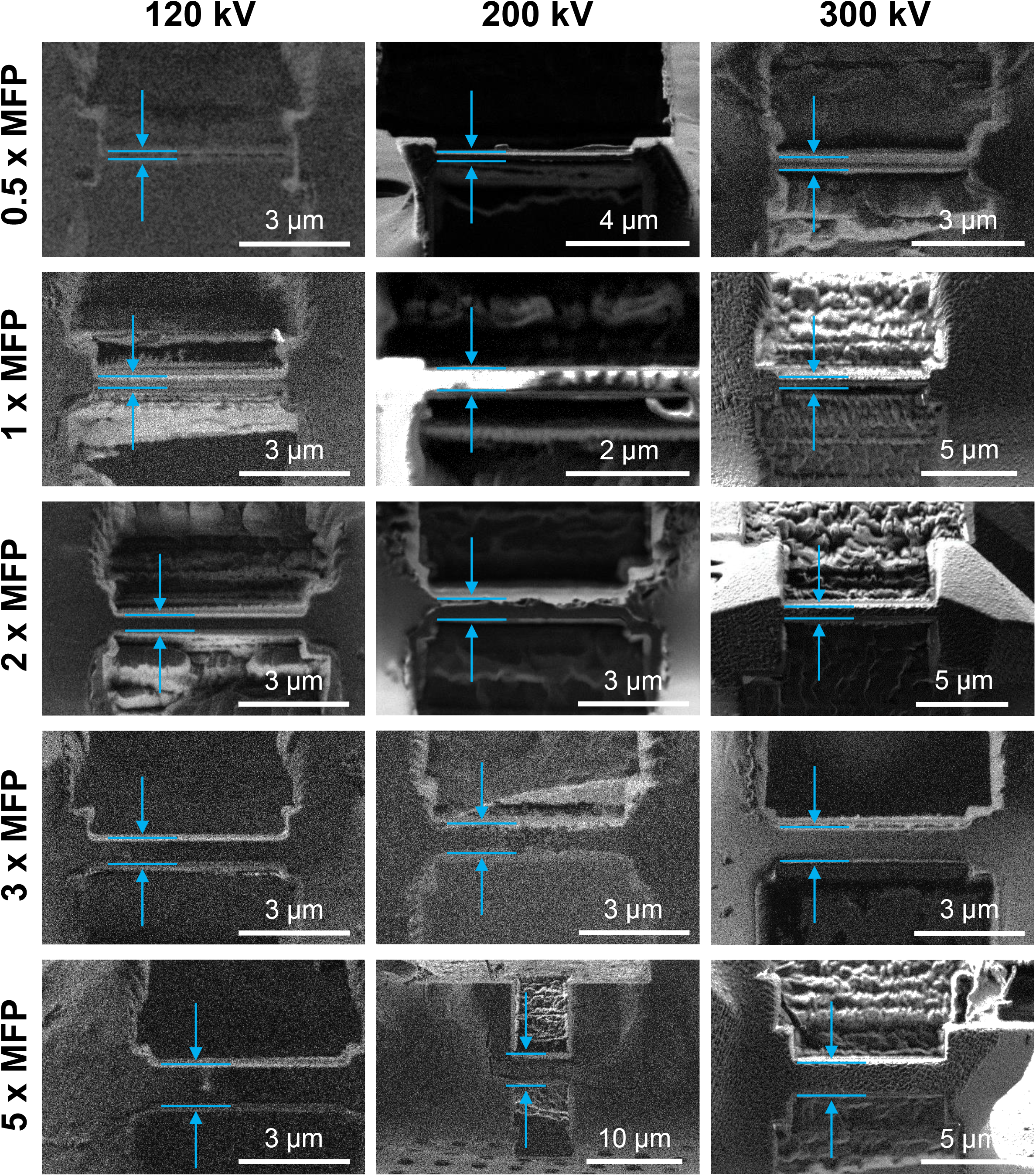
Preparing protein lamellae of variable thicknesses. Images taken by the gallium ion-beam of selected crystalline lamellae after milling. Images are sorted into rows and columns by the accelerating voltages used for data collection, and the calculated multiple of the inelastic mean free path for protein for that condition. Approximate location and size of lamellae are indicated by blue arrows and blue lines.

*MicroED experiments on crystalline lamellae*.

Each grid was carefully rotated by 90° when loading into the TEM, such that the rotation axis in the TEM was perpendicular to the milling direction in the FIB/SEM. Samples were investigated at an accelerating voltage of either 120, 200, or 300 kV. Lamellae on each grid were identified by low-magnification montaging, and the eucentric height was adjusted individually for each site. A selected area aperture was used to isolate the diffraction from a circular area approximately 3 µm in diameter from the center of each lamella. In this way, no diffraction or signal from anything other than the flat, thickness-controlled lamellae would be recorded during data collection. Continuous rotation MicroED data were collected from the real space wedge between 30° and -30° from each lamella (Figure 3).

**Figure 3.**
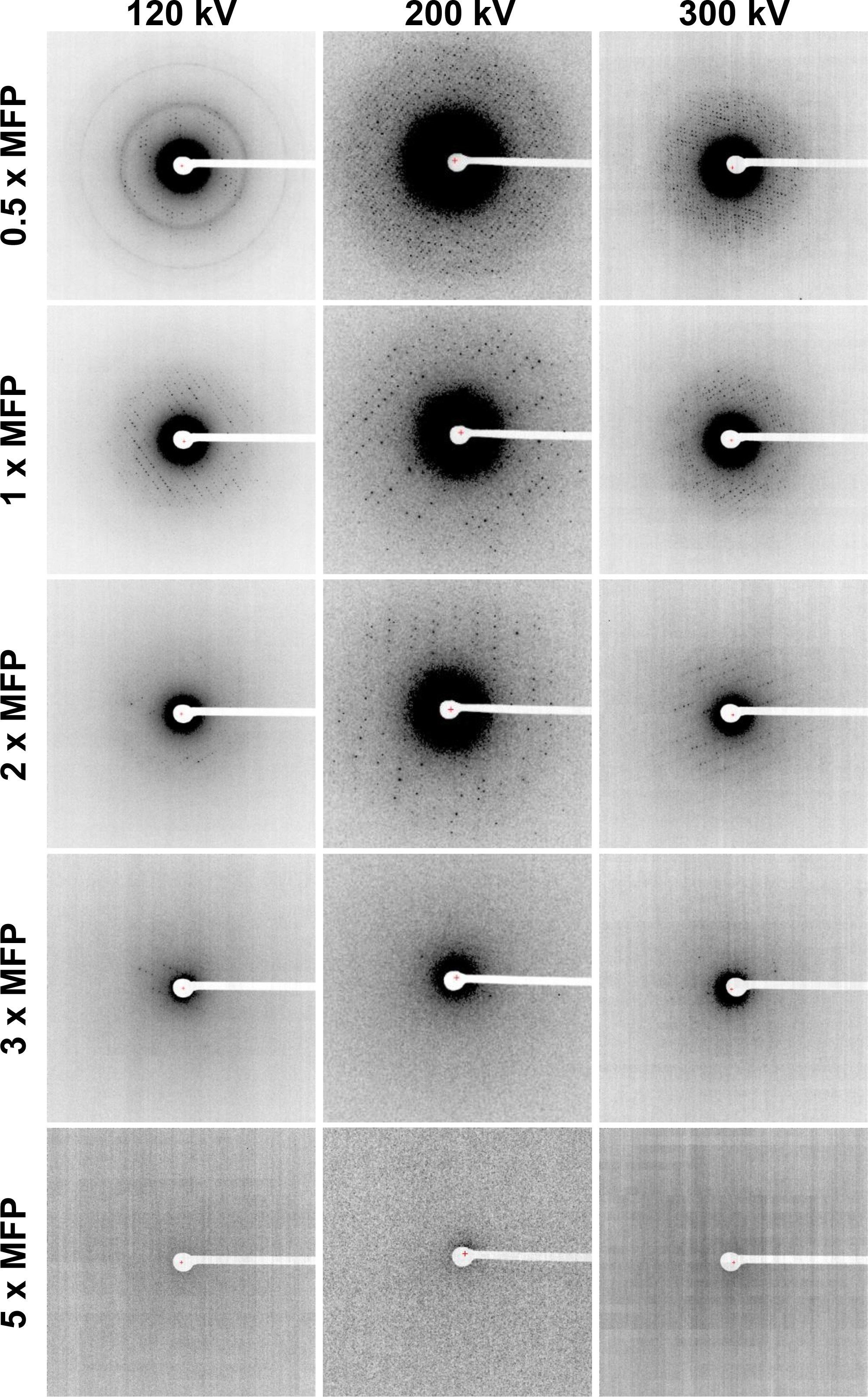
MicroED data from lamellae of different thicknesses and accelerating voltages. Single frames from MicroED datasets corresponding to 0 ° tilt. Frames are sorted into columns by the accelerating voltage and rows for the approximate thickness expressed as a multiple of the calculated inelastic mean free path at that accelerating voltage.

An estimate of resolution for a complete dataset can be obtained by visually inspecting the projection of maximum intensities through an entire continuous rotation dataset. We calculated these projections for all the lamellae (Supplementary Figures 1 – 36). These estimates corroborate the general trends seen in the integration statistics: high quality signal clearly persists to 2× MFP, greatly diminished by 3× MFP, and completely lost at approximately 4× the MFP and beyond. Measurements at 5× MFP were also taken and found to be similarly void of diffraction suggesting that at these thicknesses electrons are fully absorbed by the samples (SI Figures 24, 36).

Data from each lamella were converted to crystallographic format (Figure 3, Supplementary Figures 1 - 36). These datasets were then indexed, integrated, and scaled individually as described (Materials and Methods) (31). A resolution cutoff was applied to each dataset where the CC_1/2_ value fell to ∼33% (Table 1)(32). Integration statistics are reported for each lamella in Table 1. Collectively, strong and sharp reflections were observed from crystal lamellae that were up to 2× MFP thick at the three acceleration voltages. Little or poor diffraction was observed at 3× MFP while above 4×MFP no diffraction could be observed (Figure 3). Lamellae that yielded usable data were individually integrated and their respective models refined (Table 1, Supplementary Tables 1). In all cases, the calculated maps and composite omit maps were of high quality (Figure 4, Supplementary Figure 37).

**Figure 4.**
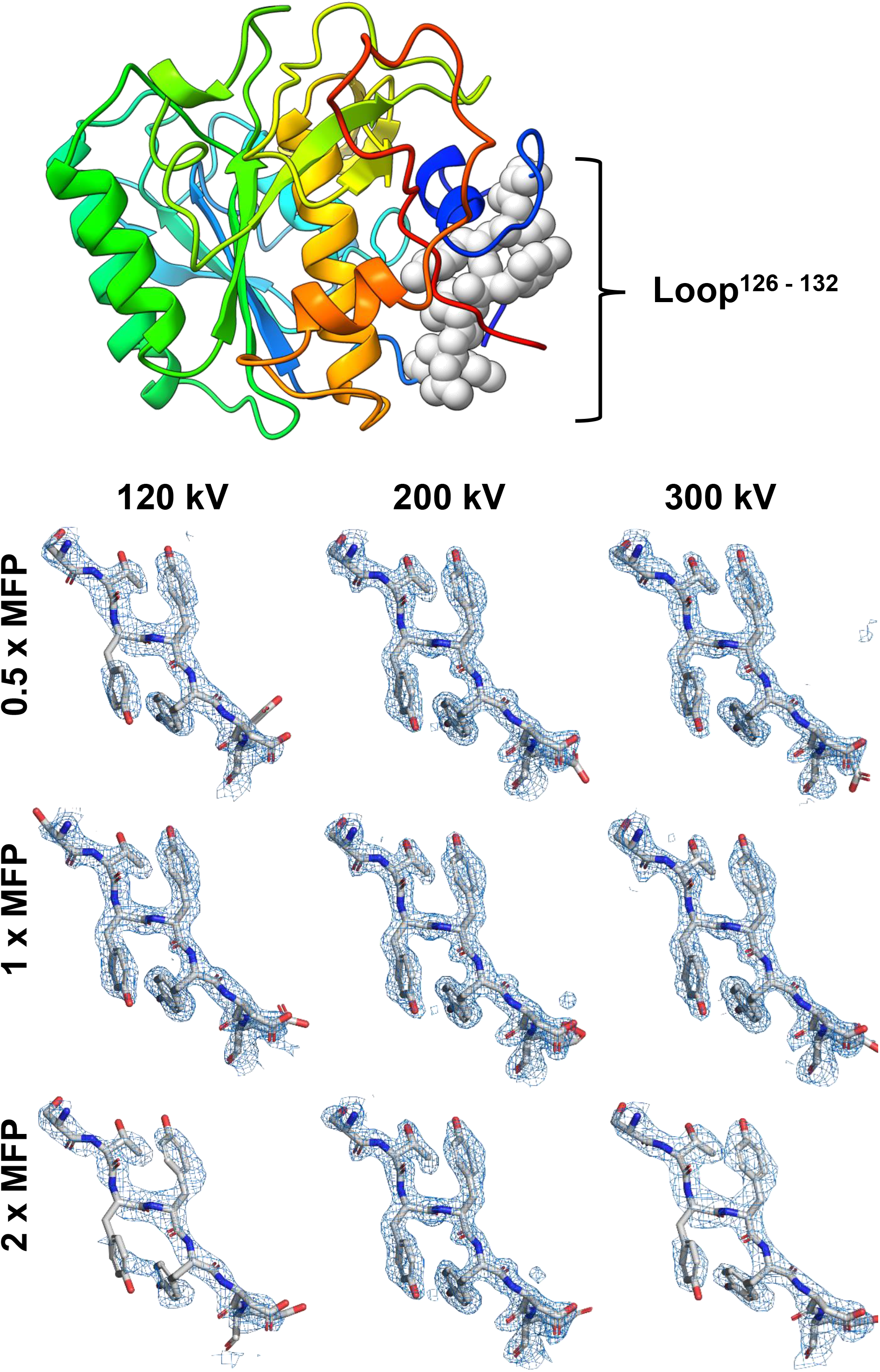
MicroED structures determined from lamellae of specified thicknesses. (Top) Final structure solution of Proteinase K in rainbow with a loop corresponding to residues 126-132 shown as gray spheres. (Bottom) 2mF_o_ -F_c_ maps from lamellae of different thicknesses and resolution cutoffs for the selected loop from above. All maps contoured at 1.5 σ level with a 2 Å carve.

**Table 1.** MicroED data from lamellae of different thicknesses

|  |  |  |  |  |  |  |  |  |  |  |
| --- | --- | --- | --- | --- | --- | --- | --- | --- | --- | --- |
| 120 kV | Crystal # | 1 | 2 | 3 | 4 | 5 |  |  |  |  |
|  | Thickness (nm) | 130 | 200 | 325 | 600 | 960 |  |  |  |  |
|  | MFP (x) | 0.6 | 0.9 | 1.5 | 2.8 | 4.5 |  |  |  |  |
|  | Resolution limit (Å) | 2.3 | 2.0 | 2.7 | 3.8* | 20** |  |  |  |  |
|  | Completeness (%) | 82.5 | 87.7 | 82.6 | - | - |  |  |  |  |
|  | R-pim (%) | 14.9 | 12.8 | 29.6 | - | - |  |  |  |  |
|  | <I / σ(I)> | 5.0 | 4.9 | 3.2 | 0*** | 0*** |  |  |  |  |
|  | CC<sub>1/2</sub> | 97.5 | 98.3 | 89.3 | 0*** | 0*** |  |  |  |  |
|  | R work (%) | 22.2 | 21.5 | 24.1 | - | - |  |  |  |  |
|  | R free (%) | 26.1 | 24.3 | 28.5 | - | - |  |  |  |  |
| 200 kV | Crystal # | 1 | 2 | 3 | 4 | 5 | 6 | 7 | 8 | 9 |
|  | Thickness (nm) | 95 | 115 | 130 | 260 | 460 | 530 | 540 | 800 | 1400 |
|  | MFP (x) | 0.3 | 0.4 | 0.5 | 1 | 1.7 | 1.95 | 2 | 2.9 | 5.1 |
|  | Resolution limit (Å) | 2.35 | 1.85 | 1.95 | 1.95 | 1.95 | 2.3 | 2.4 | 4.1* | 20** |
|  | Completeness (%) | 86.6 | 86.0 | 96.8 | 91.1 | 91.9 | 90.0 | 78.6 | - | - |
|  | R-pim (%) | 15.9 | 10.8 | 13.4 | 13.3 | 15.4 | 19.7 | 15.6 | - | - |
|  | <I / σ(I)> | 3.6 | 5.2 | 4.4 | 4.25 | 3.85 | 4.41 | 3.25 | - | - |
|  | CC1/2 (%) | 96.8 | 98.8 | 97.9 | 97.6 | 96.3 | 91.7 | 92.5 | 0*** | 0*** |
|  | R work (%) | 20.0 | 18.1 | 19.4 | 18.7 | 20.1 | 20.1 | 19.3 | - | - |
|  | R free (%) | 24.0 | 21.7 | 23.6 | 23.9 | 24.5 | 24.1 | 22.2 | - | - |
| 300 kV | Crystal # | 1 | 2 | 3 | 4 | 5 | 6 | 7 |  |  |
|  | Thickness (nm) | 150 | 170 | 320 | 360 | 550 | 880 | 1650 |  |  |
|  | MFP (x) | 0.47 | 0.54 | 1.0 | 1.2 | 1.7 | 2.8 | 5.2 |  |  |
|  | Resolution limit (Å) | 1.9 | 2.1 | 2.1 | 2.05 | 2.9 | 3.7* | 20** |  |  |
|  | Completeness (%) | 90.5 | 89.9 | 82.5 | 92.9 | 90 | - | - |  |  |
|  | R-pim (%) | 11.8 | 32.4 | 14.4 | 12.8 | 23.1 | - | - |  |  |
|  | <I / σ(I)> | 4.9 | 3.38 | 3.61 | 3.8 | 2.43 | - | - |  |  |
|  | CC1/2 (%) | 98.5 | 91.7 | 96.4 | 95.6 | 77.3 | 0*** | 0*** |  |  |
|  | R work (%) | 19.17 | 27.25 | 19.85 | 19.14 | 23.8 | - | - |  |  |
|  | R free (%) | 22.64 | 33.20 | 24.46 | 23.64 | 27.45 | - | - |  |  |

\*Value approximated from diffraction images that could not be automatically processed
\*\*Best possible value taken from beamstop cut off
\*\*\*Value presumed from inability to integrate datasets

## Discussion

The relationship between crystal thickness, acceleration voltage, and the quality of MicroED data and the subsequently determined structures was systematically investigated. The samples used here were of the same protein, grown in the same batch, from crystals of originally similar size, and machined into lamellae using the same protocol (Materials and Methods). MicroED data were collected using established procedures (17, 23, 33). The data were collected at the three most common acceleration voltages used in cryo-EM - 120, 200, and 300kV. No increasing, systematic errors prevented structure determination from crystals up to twice the MFP. Even thinner crystals did not appear to have any significant advantage (Figure 5, Table 1). Indeed, the crystallographic statistics, metrics of the determined structures, and resulting maps from lamellae up to 2× MFP all appear relatively similar (Table 1 and Figure 5). However, beyond 2×MFP the overall resolution and quality metrics rapidly fall off (Figure 5). No structures could be determined at 3×MFP for all acceleration voltages, although sporadic diffraction spots at low resolution were visible on some images. No diffraction was observed for samples thicker than 3×MFP, suggesting that most electrons are absorbed by the sample. These observations are in agreement with the measurements of single diffraction patterns from Zhou et al. (24), where the best data were observed at intermediate thicknesses rather than at an extremum.

**Figure 5.**
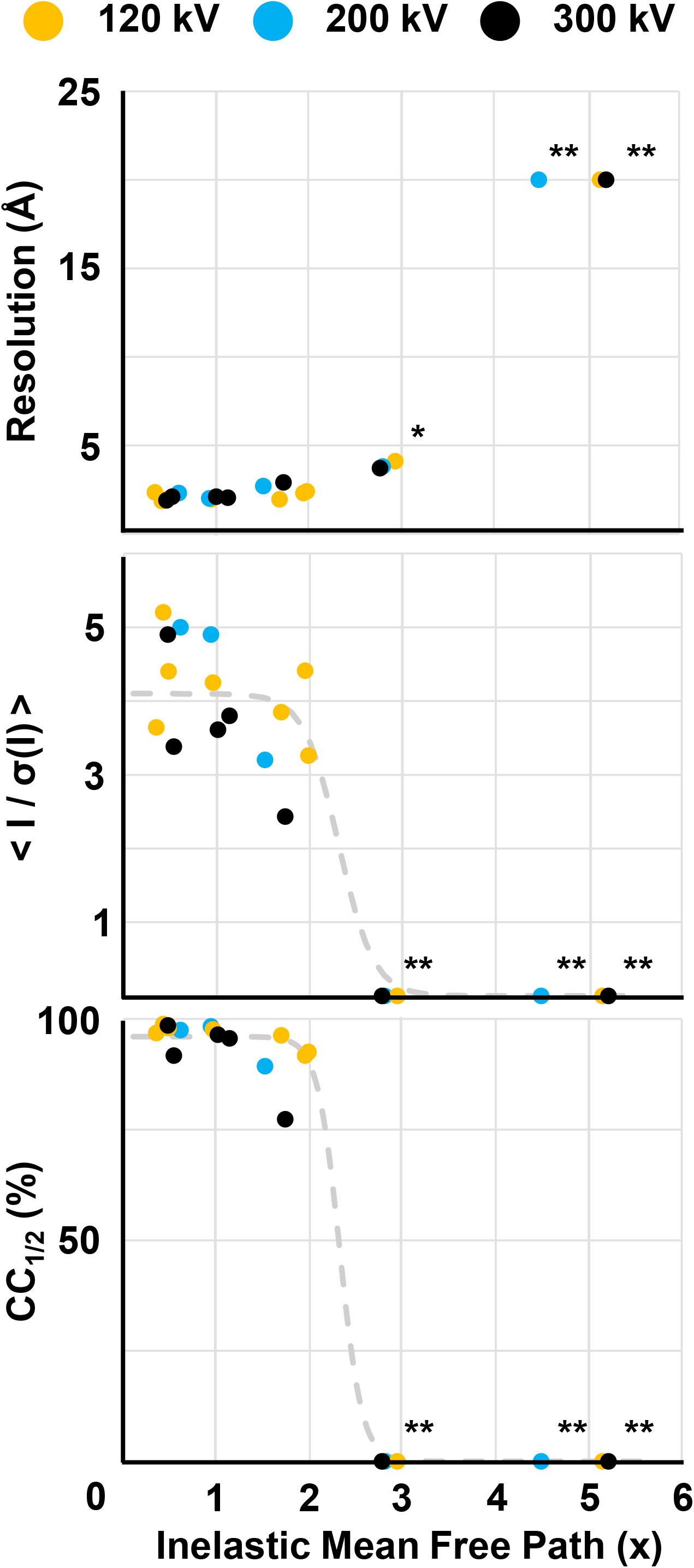
Data quality metrics as a function of thickness. (A) Resolution cutoff, (B) mean signal to noise, and (C) the half set correlation coefficient for all the measured crystals. Data points are color coded according to accelerating voltage as indicated. A simple reverse sigmoid function is fit to the data in (B) and (C) to demonstrate the sharp drop in quality after approximately 2× MFP. (*) Value approximated from diffraction images that could not be automatically processed. (**) Best possible value given experimental cutoff due to beamstop. (***) Value presumed from inability to integrate datasets

The data suggests that the largest detriment to determining structures by MicroED from thicker samples appears to be absorption. Reliable high quality structures can be determined from up to 2× MFP regardless of acceleration voltage. Beyond that point, the resolution deteriorated and structures became difficult to determine. Absorption effects are compounded by the fact that thicker crystals produce more diffuse scattering, and increased inelastic scattering results in higher background noise. Taken together, it is clear that the elastic, Bragg peaks are diminished by absorption in thick samples, and that the higher resolution reflections are quickly lost to additional noise from the increased background. It is possible that an energy filter could help mitigate some of these issues, and we expect that the usable thickness could slightly increase with this addition. Initial results already demonstrated that an energy filter leads to an increase in the signal to noise ratio and the attainable resolution (34–38).

Computer simulations are at odds with MicroED experimental results. All simulations are inherently limited by the validity of their assumptions, as correctly discussed in Subramanian *et al.* (10). We suggest that at least five assumptions are inadequate in current electron diffraction simulations. Namely most assume that: 1. macromolecular crystals are perfect; 2. diffraction is only collected from a major zone axis (always true for 2D crystals, but rarely for 3D crystals); 3. contribution from disordered solvent is negligible 4. data are collected from stationary crystals (MicroED uses continuous rotation) and 5. inelastic scattering, or absorption is not significant. Though more recent simulations have made progress in accounting for some of these discrepancies (39), future simulations would benefit from a more accurate modelling of the experimental setup.

We demonstrated that high quality structures can be obtained from samples that are up to 2× the MFP regardless of the acceleration voltage. This corresponds to a real space thicknesses of approximately 430, 540, and 640 nm for the accelerating voltages of 120, 200, and 300 kV, respectively. We find that samples that were thicker did not yield usable data, and electrons were generally lost to absorption at thicknesses of 4× MFP or more. We expect that these limits could be somewhat relaxed with an energy filter, but the exact parameters will need to be investigated in future studies. Importantly, as FIB milling becomes a standard method for sample preparation for macromolecular MicroED studies, aiming for a thicknesses of less than 2×MFP will maximize the likelihood of producing the best data and highest quality structures. Regardless of the cryo-EM method employed, the current study provides a benchmark for the sample thickness in cryo-EM especially for electron tomographic studies of FIB milled tissues and cells.

## Acknowledgements

This study was supported by the National Institutes of Health P41GM136508. The Gonen laboratory is supported by funds from the Howard Hughes Medical Institute.

## Materials and Methods

### Materials

Proteinase K (*E. Album*) and ammonium sulfate were purchased from Sigma (Sigma-Aldrich, St. Louis, MO) and used without further purification. Stock solutions were made using Milli-Q water filtered three times through a 0.4 µm porous membrane.

### Crystallization

Proteinase K was crystallized in batch by dissolving 1 mg of lyophilized protein powder with 200 µL of 2M ammonium sulfate at room temperature. 10-30 µm crystals formed within 10 minutes.

### Grid preparation

Quantifoil Cu 200 R 2/2 grids were glow discharged for 30 s immediately prior to use. Grids were vitrified using a Leica GP2 vitrification robot at room temperature. The sample chamber was set to 95% relative humidity and the filter paper equilibrated in the humid air for 15 min prior to grid preparation. The tube of crystals was gently shaken just before 3 µL of protein crystal solution was removed and gently pipetted onto the carbon side of the grid in the vitrification chamber. The slurry was incubated on the grid for 30 s. The grid was then gently blotted from the back for 20 s, plunged into liquid ethane, and transferred to liquid nitrogen for storage.

### Focused ion-beam and scanning electron microscopy

Vitrified grids were transferred into a cryogenically cooled Thermo-Fisher Aquilos dual beam FIB/SEM. The grids were coated in a thin layer of fine platinum followed by a thick >100 nm layer of coarse platinum by sputter coating to protect the crystals from the damaging ion and electron beams (29). Whole grid atlases were recorded using the MAPS software (Thermo-Fisher), where individual crystals were identified and aligned to eucentric height. Twenty-one crystals over six grids were chosen for milling. Crystals were milled as described (23, 29, 30, 40, 41). Briefly, the milling was conducted in steps of: rough milling, fine milling, and polishing. Each step was performed on each crystal sequentially prior to advancing to the next step to reduce the effects of amorphous ice buildup and contamination. Rough milling steps were conducted at 100 pA and removed all but 5 µm of crystalline material. Fine milling used an ion-beam current of 50 pA and removed material up to 250 nm away from the desired thickness. Polishing was conducted at an ion-beam current of 10 pA and was used until the approximate final thickness was achieved. A final image of the lamella was taken using the ion-beam at 1.5 pA (Figure 2, Supplementary Figures 1 – 36) to assess the final thickness using the measurement tool in the Aquilos user interface. All micrographs taken by the scanning electron beam were performed at an accelerating voltage of 5 kV and a beam current of 1.6 pA. All ion-beam imaging and milling was conducted at an accelerating voltage of 30 kV.

### MicroED data collection

Grids containing milled lamellae were transferred to either a Thermo-Fisher Titan Krios G3i or Talos Arctica transmission electron microscopes. The Krios was operated at an accelerating voltage of either 120 or 300 kV, whereas the Arctica operated at an accelerating voltage of 200 kV. Electrons at the accelerating voltages of 120, 200 or 300 kV have corresponding de Broglie wavelengths of 0.0335 Å, 0.0251 Å, or 0.0197 Å, respectively. Both microscopes were equipped with a field emission gun and a Ceta-D 16M (4k×4k) CMOS detector. Lamellae were identified on each grid by taking a low-magnification montage, where they appeared as thin white stripes against an otherwise dark background. Crystalline lamellae within these strips appeared semi-transparent and suspended over this gap. Lamellae were brought to eucentric height and initially evaluated by taking a single 1 s exposure in diffraction mode at 0° stage tilt. MicroED data were collected as described (33). In short, the stage was continuously rotated at a rate of 0.25° s^-1^ while the crystal was illuminated in a parallel electron beam. Frames were read out from the detector every 1 or 2 seconds and saved as a stack in MRC format. Each dataset corresponded to 120 - 240 images corresponding to the real-space angular wedge between +30 to -30°. The electron beam was approximately 10 µm in diameter, where the corresponding exposure was calibrated to a rate of approximately 0.01 e^-^ Å^-2^ s^-1^. The total exposure to each lamellae was therefore approximately 2.4 e^-^ Å^-2^. Signal from the center of each lamella was isolated by inserting a selected area aperture of 100 µm for the Arctica or 150 µm for the Krios, corresponding to an area of approximately 3 µm in diameter projected from the sample for either microscope.

### MicroED data processing

MRC stacks were converted to individual frames in SMV format using the MicroED tools as described (23, 33). MicroED tools can be downloaded freely at (https://cryoem.ucla.edu/downloads). Reflections were indexed and integrated in XDS (31). Individual or groups of datasets were scaled in either AIMLESS (42) or XSCALE (43). Each dataset was used to determine the structure using molecular replacement in Phaser with the search model PDB 6CL7 (44, 45). Models were refined in phenix.refine (46) using electron scattering factors.

### Calculations

The inelastic mean free path (MFP) was calculated using the original formulation by Malis *et al.* (6) and Egerton (47) et al. and subsequently used by Feja et al. (48) and Grimm et al. (8) given by:

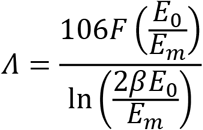

and

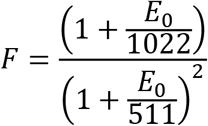

where

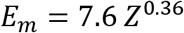

We similarly used the values of β = 10 mrad, Z = 8, and E_0_ for the acceleration voltage used, e.g. 120, 200, or 300 kV as the typical values employed in those investigations.

An optical refractive term was applied as suggested in Grimm *et al*. (8), and a value of n = 1.48 was chosen based on the determined value for lysozyme layers (49). This translates to inelastic mean free path values of 214, 272, and 317 nm for 120, 200, and 300 kV accelerating voltages. These values are in good agreement with those previously measured experimentally (6–8, 47, 48, 50) and have a direct relationship to the deposited dose (51).

The mean-free path is closely related to the collision stopping power, as calculated using e.g. ESTAR (52). For a typical protein sample with density 1.17 g cm^-3^ (53), the tabulated stopping power implies that a 120 kV electron loses 4.15 MeV per cm of traversed sample (3.22 MeV and 2.72 MeV for 200 kV and 300 kV electrons, respectively). The deposited energy per unit length calculated by this method was previously used to derive estimates of the dose from a given exposure (45, 51) and its inverse is correlated to the mean-free path length used here with an asymptotic standard error of <2%.

The curves presented in Figure 5 (B) and (C) are reverse sigmoid functions of the form:

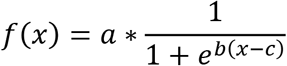

The functions were fit using simple least squares.

## Supplementary Appendix

**Supplementary Figure 1.**
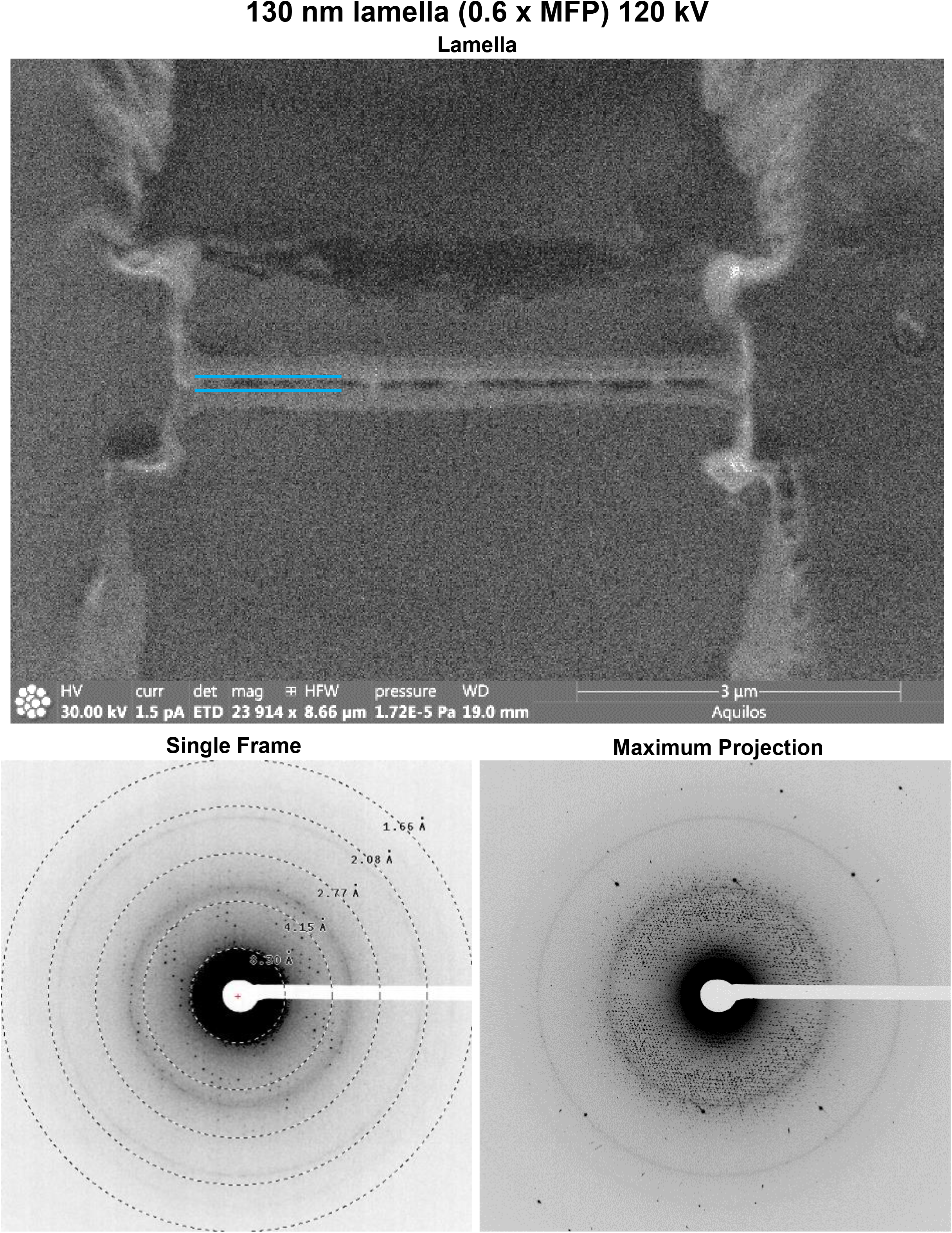
Data from a 130 nm lamella of proteinase K collected at 120 kV. (Top) Image in the FIB directly after the milling finished. (Bottom Left) Single frame of MicroED data (Bottom Right) Maximum projection of the entire MicroED dataset onto a single frame.

**Supplementary Figure 2.**
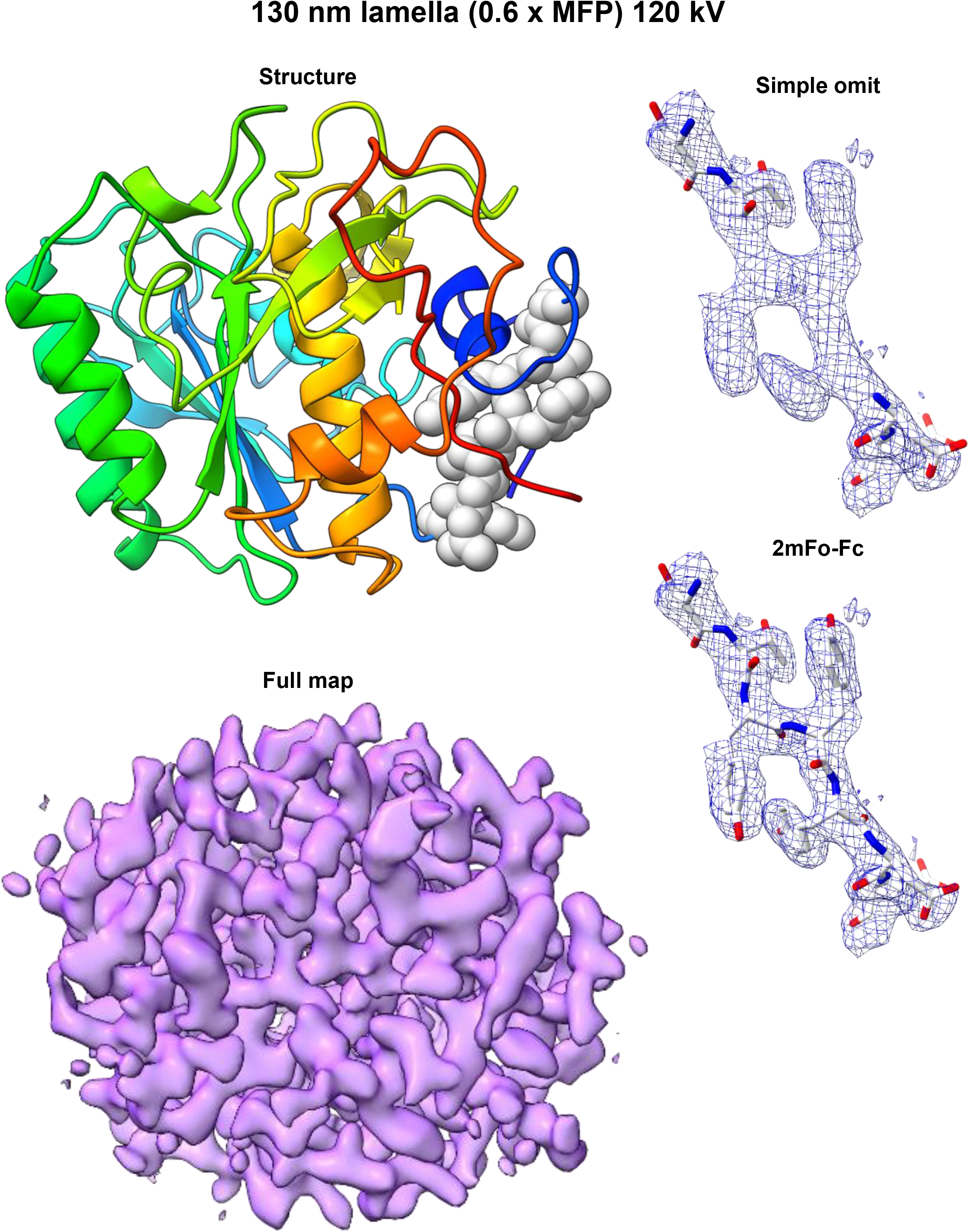
Structure of Proteinase K determined from a 130 nm lamella at 120 kV. (Left Top) Cartoon structure. (Left Bottom) 2mF_o_-F_c_ map contoured at 1.5σ level. (Right Top) 2mF_o_-F_c_ map of the structure generated without three loop residues indicated in the structure on the right. (Right bottom) 2mF_o_-F_c_ map for the same region calculated with the missing residues.

**Supplementary Figure 3.**
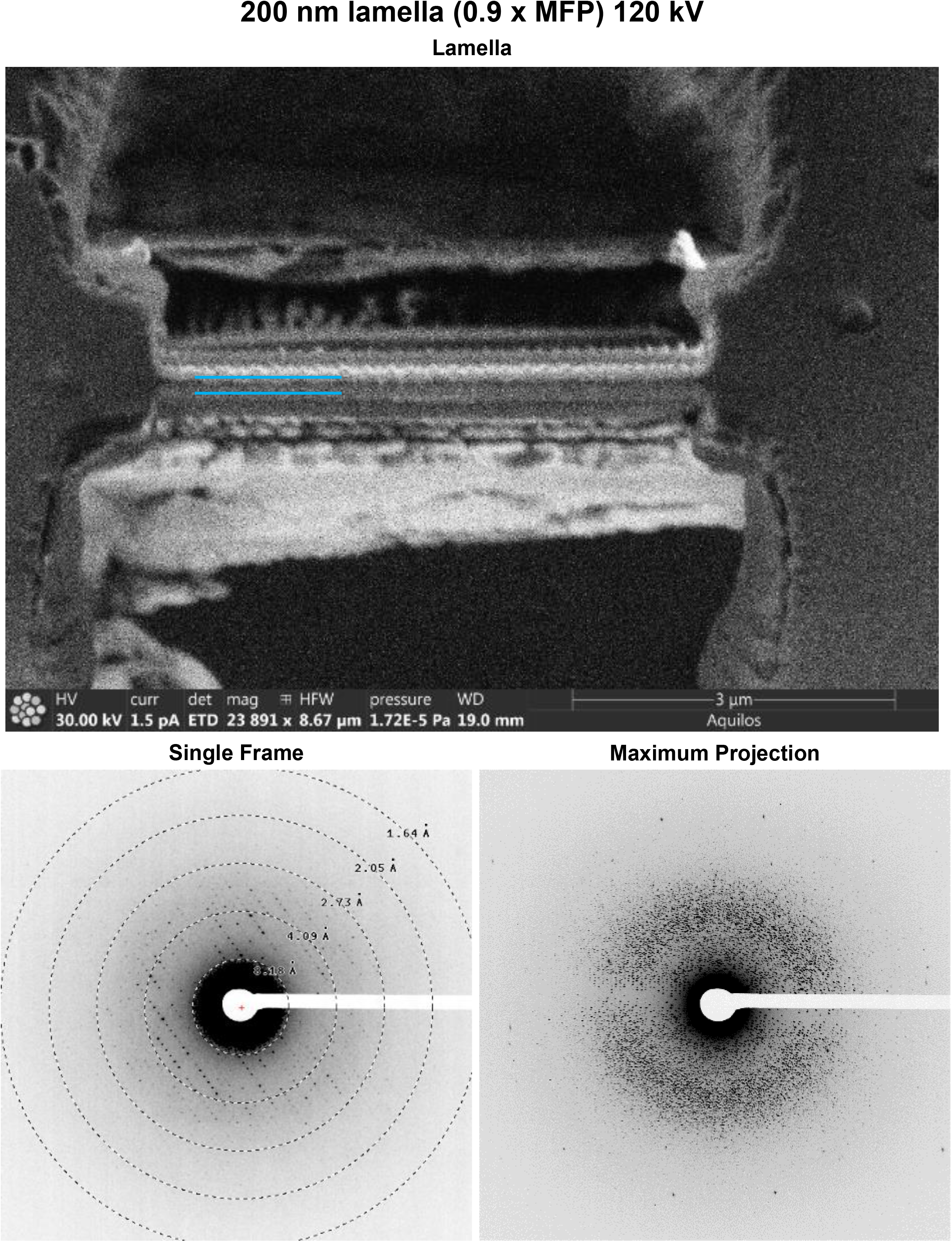
Data from a 200 nm lamella of proteinase K collected at 120 kV. (Top) Image in the FIB directly after the milling finished. (Bottom Left) Single frame of MicroED data (Bottom Right) Maximum projection of the entire MicroED dataset onto a single frame.

**Supplementary Figure 4.**
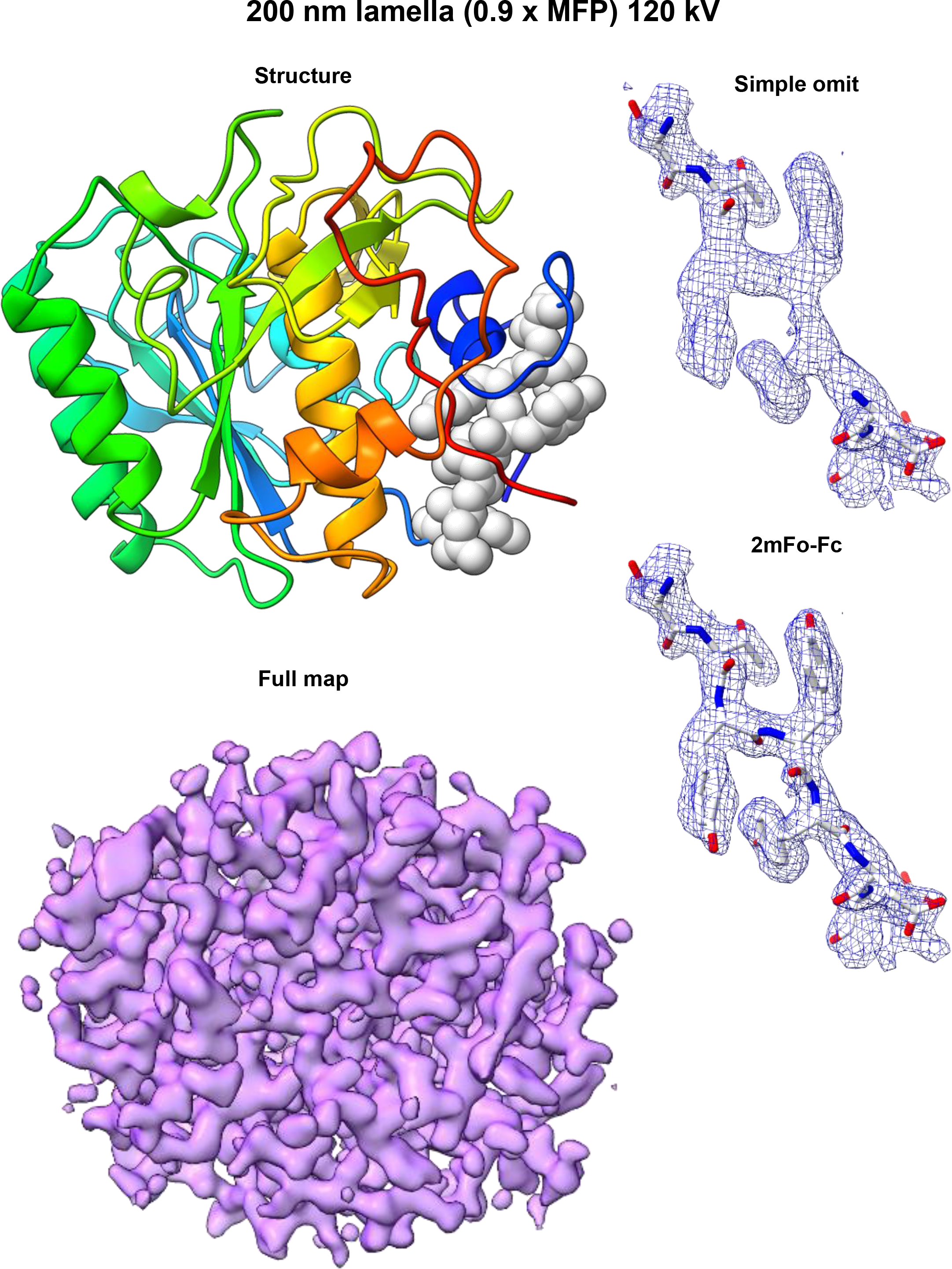
Structure of Proteinase K determined from a 200 nm lamella at 120 kV. (Left Top) Cartoon structure. (Left Bottom) 2mF_o_-F_c_ map contoured at 1.5σ level. (Right Top) 2mF_o_-F_c_ map of the structure generated without three loop residues indicated in the structure on the right. (Right bottom) 2mF_o_-F_c_ map for the same region calculated with the missing residues.

**Supplementary Figure 5.**
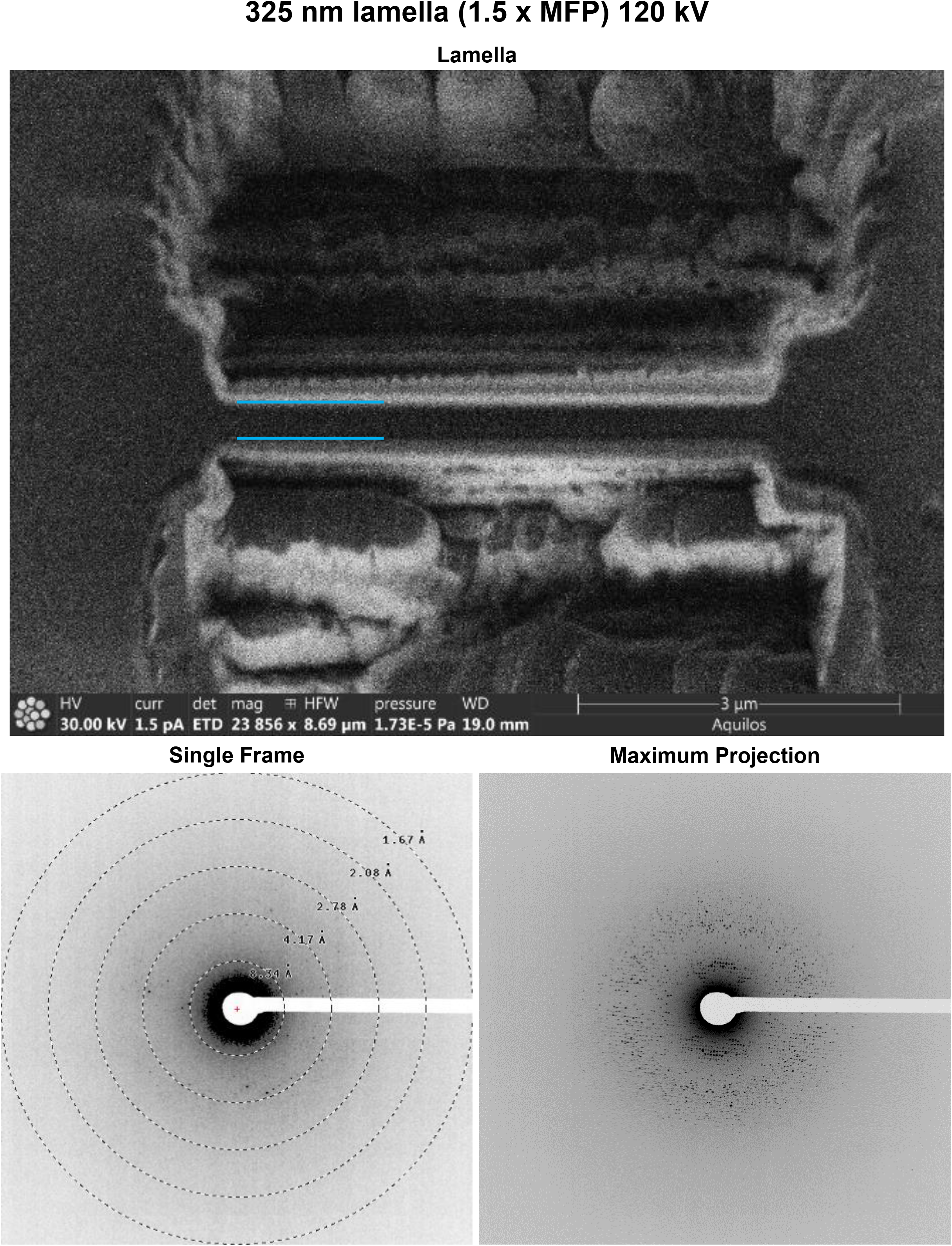
Data from a 325 nm lamella of proteinase K collected at 120 kV. (Top) Image in the FIB directly after the milling finished. (Bottom Left) Single frame of MicroED data (Bottom Right) Maximum projection of the entire MicroED dataset onto a single frame.

**Supplementary Figure 6.**
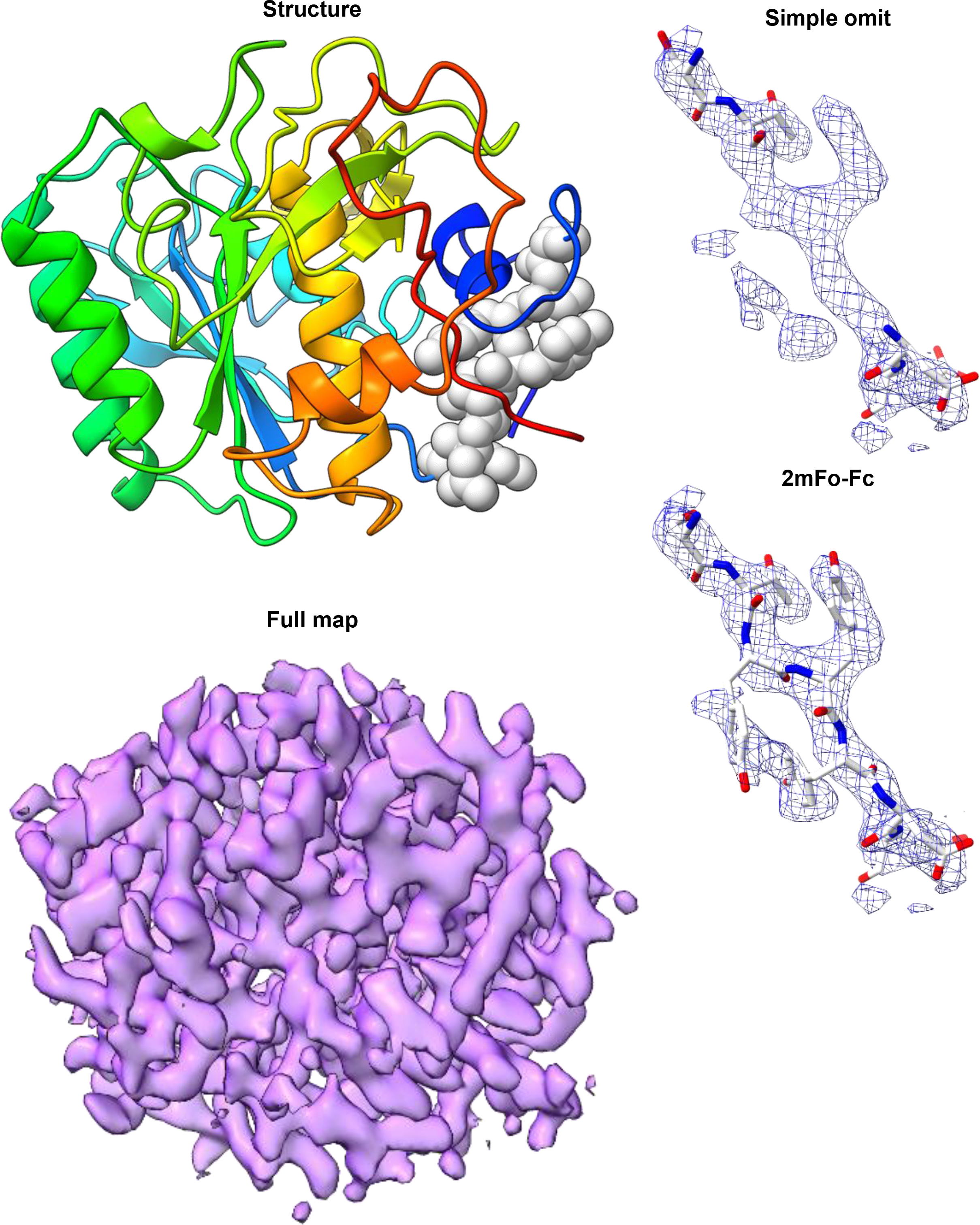
Structure of Proteinase K determined from a 325 nm lamella at 120 kV. (Left Top) Cartoon structure. (Left Bottom) 2mF_o_-F_c_ map contoured at 1.5σ level. (Right Top) 2mF_o_-F_c_ map of the structure generated without three loop residues indicated in the structure on the right. (Right bottom) 2mF_o_-F_c_ map for the same region calculated with the missing residues.

**Supplementary Figure 7.**
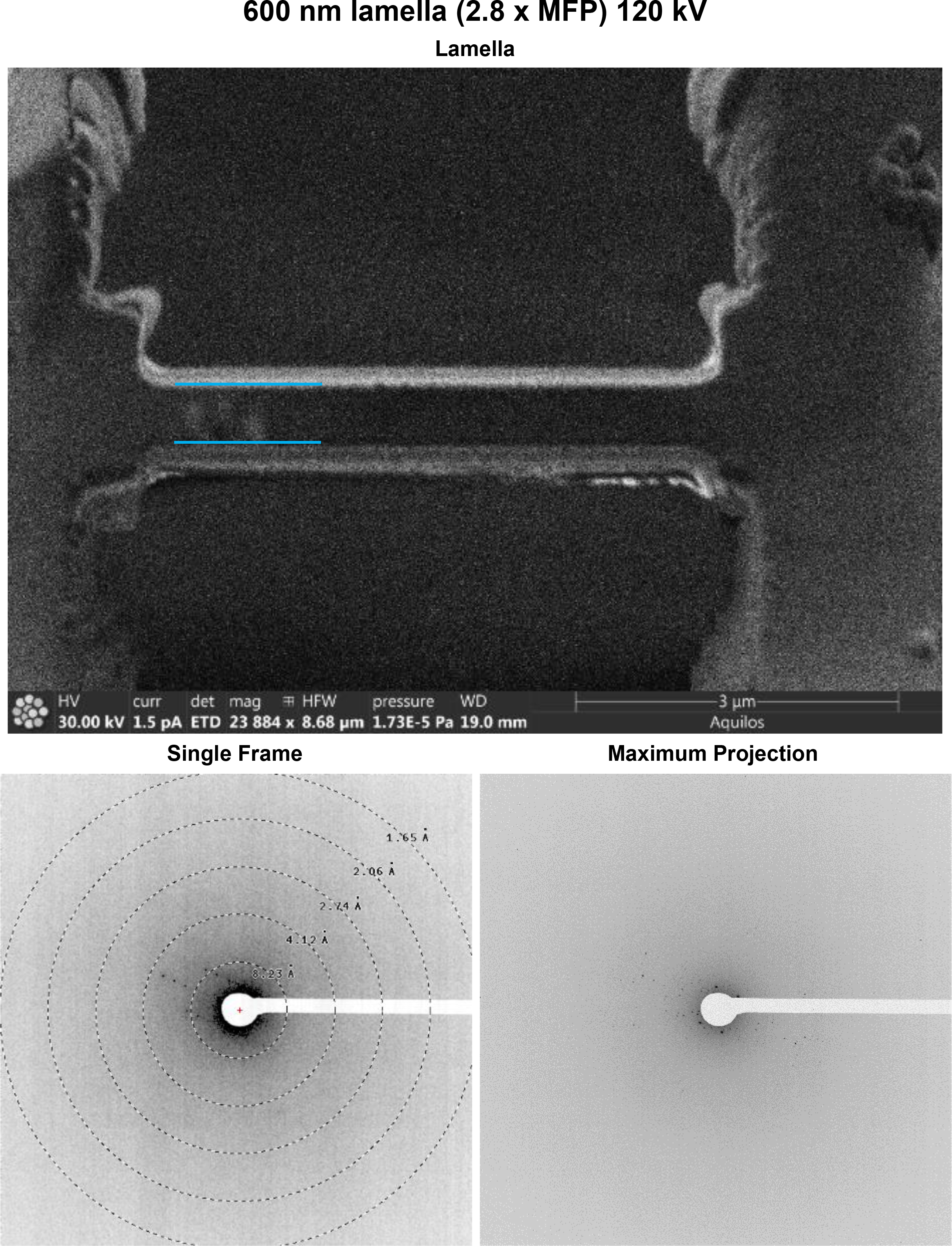
Data from a 600 nm lamella of proteinase K collected at 120 kV. (Top) Image in the FIB directly after the milling finished. (Bottom Left) Single frame of MicroED data (Bottom Right) Maximum projection of the entire MicroED dataset onto a single frame.

**Supplementary Figure 8.**
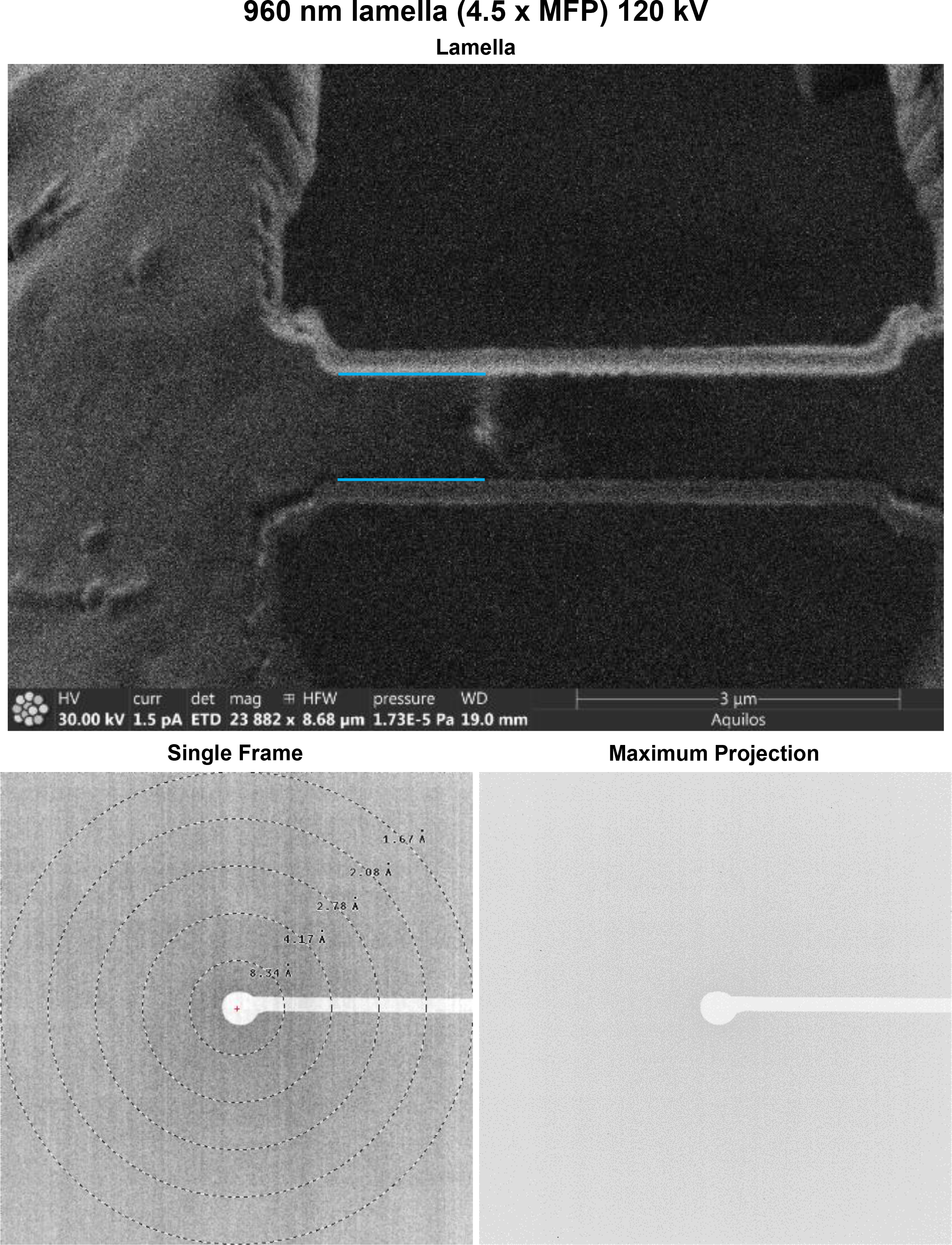
Data from a 960 nm lamella of proteinase K collected at 120 kV. (Top) Image in the FIB directly after the milling finished. (Bottom Left) Single frame of MicroED data (Bottom Right) Maximum projection of the entire MicroED dataset onto a single frame.

**Supplementary Figure 9.**
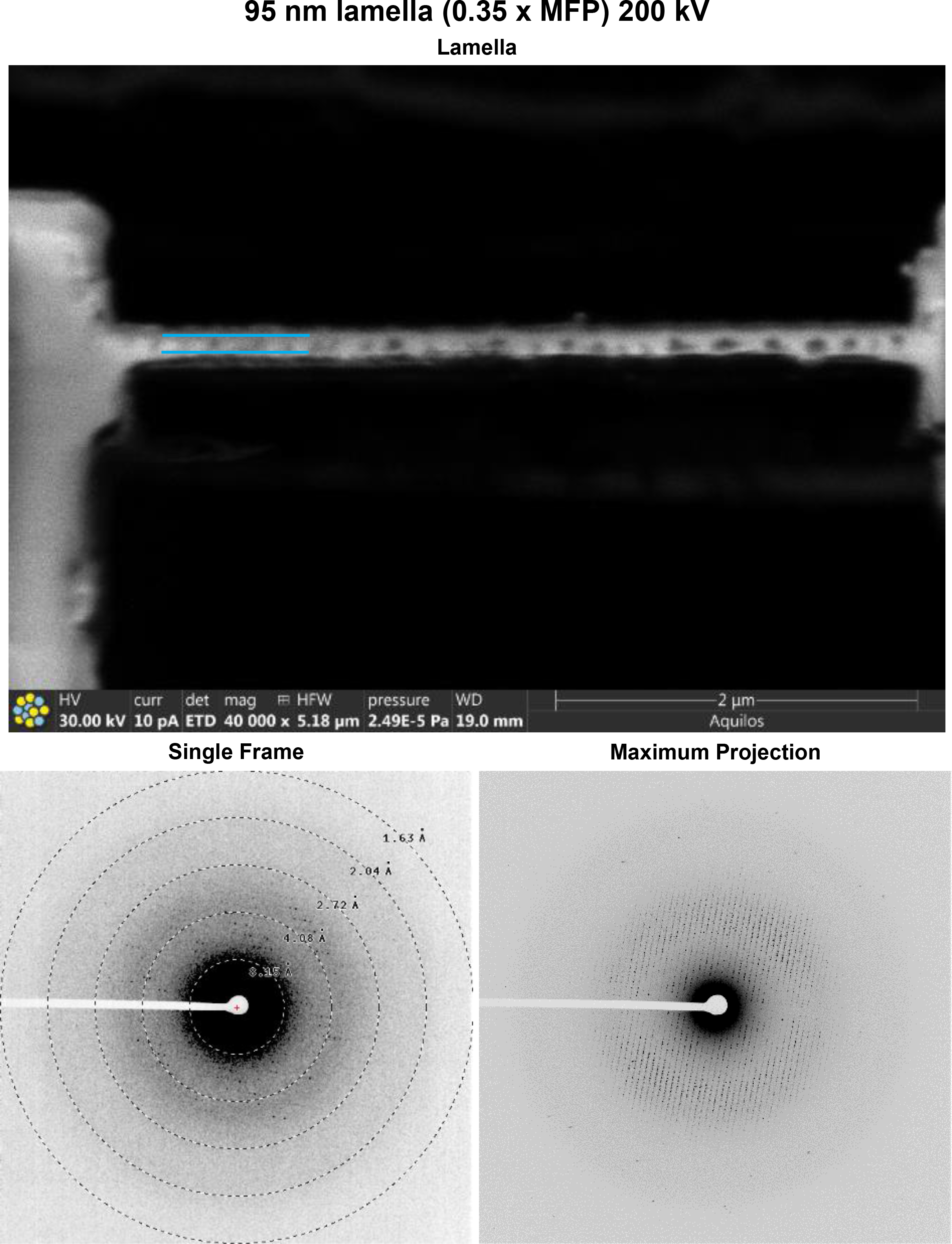
Data from a 95 nm lamella of proteinase K collected at 200 kV. (Top) Image in the FIB directly after the milling finished. (Bottom Left) Single frame of MicroED data (Bottom Right) Maximum projection of the entire MicroED dataset onto a single frame.

**Supplementary Figure 10.**
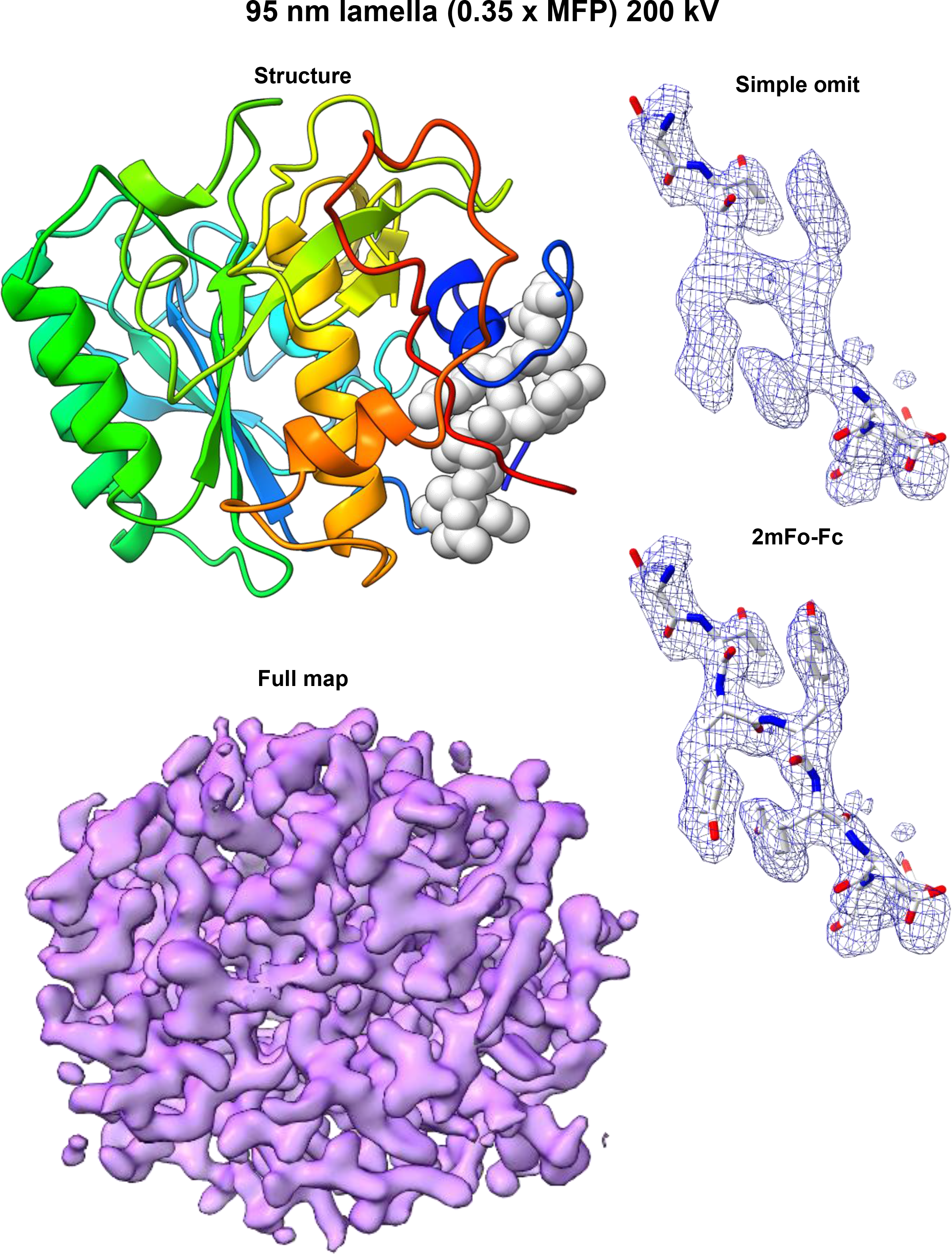
Structure of Proteinase K determined from a 95 nm lamella at 200 kV. (Left Top) Cartoon structure. (Left Bottom) 2mF_o_-F_c_ map contoured at 1.5σ level. (Right Top) 2mF_o_-F_c_ map of the structure generated without three loop residues indicated in the structure on the right. (Right bottom) 2mF_o_-F_c_ map for the same region calculated with the missing residues.

**Supplementary Figure 11.**
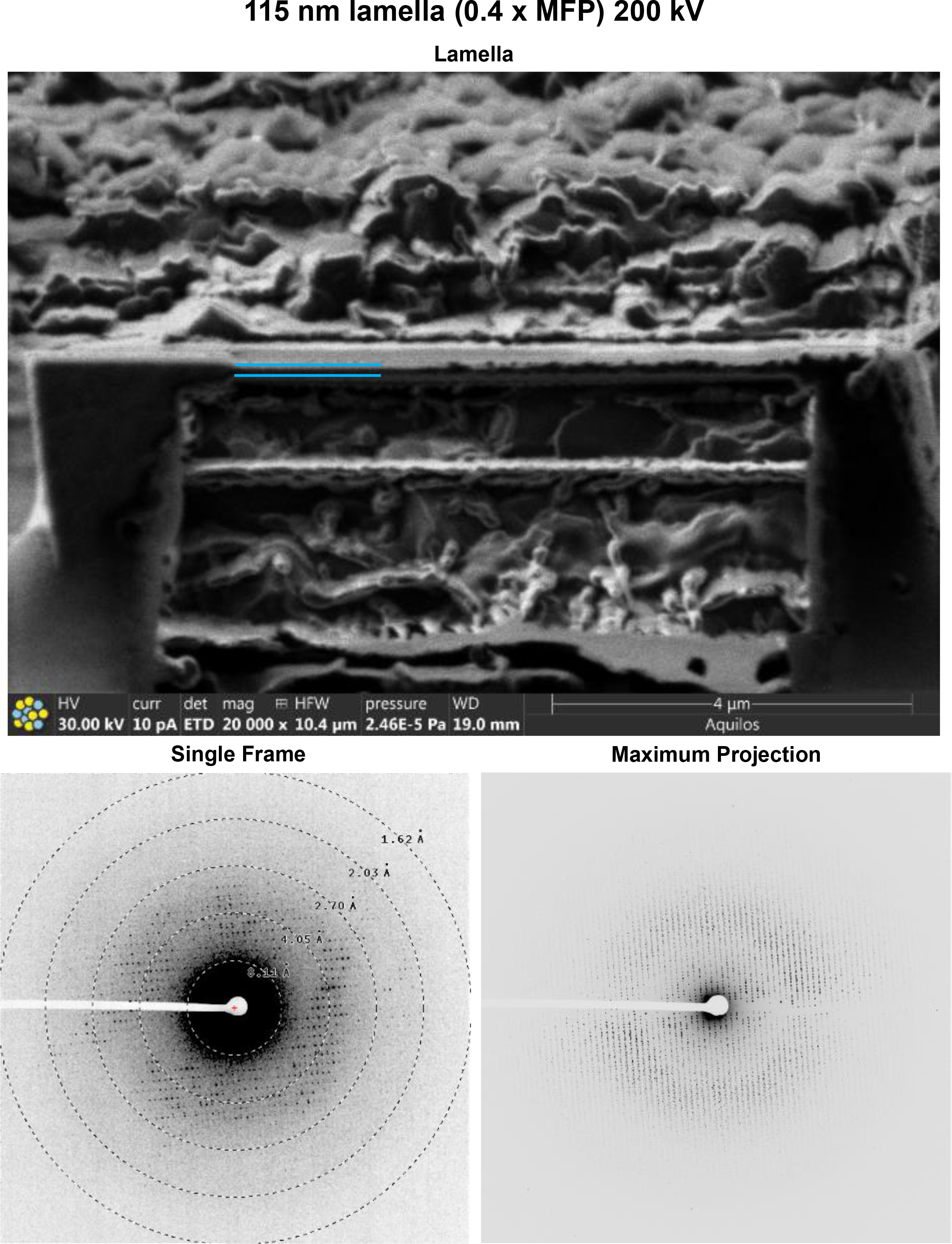
Data from a 115 nm lamella of proteinase K collected at 200 kV. (Top) Image in the FIB directly after the milling finished. (Bottom Left) Single frame of MicroED data (Bottom Right) Maximum projection of the entire MicroED dataset onto a single frame.

**Supplementary Figure 12.**
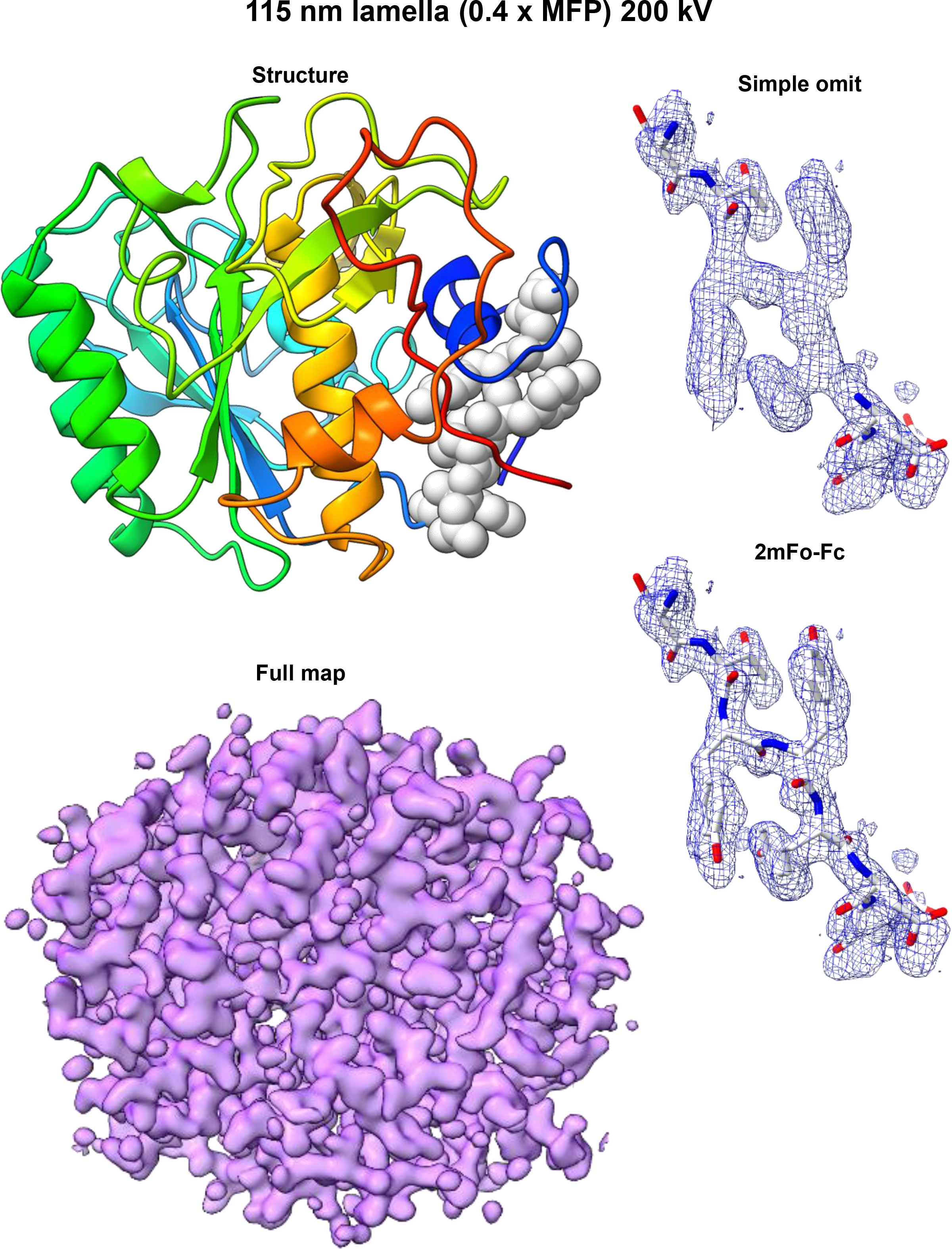
Structure of Proteinase K determined from a 115 nm lamella at 200 kV. (Left Top) Cartoon structure. (Left Bottom) 2mF_o_-F_c_ map contoured at 1.5σ level. (Right Top) 2mF_o_-F_c_ map of the structure generated without three loop residues indicated in the structure on the right. (Right bottom) 2mF_o_-F_c_ map for the same region calculated with the missing residues.

**Supplementary Figure 13.**
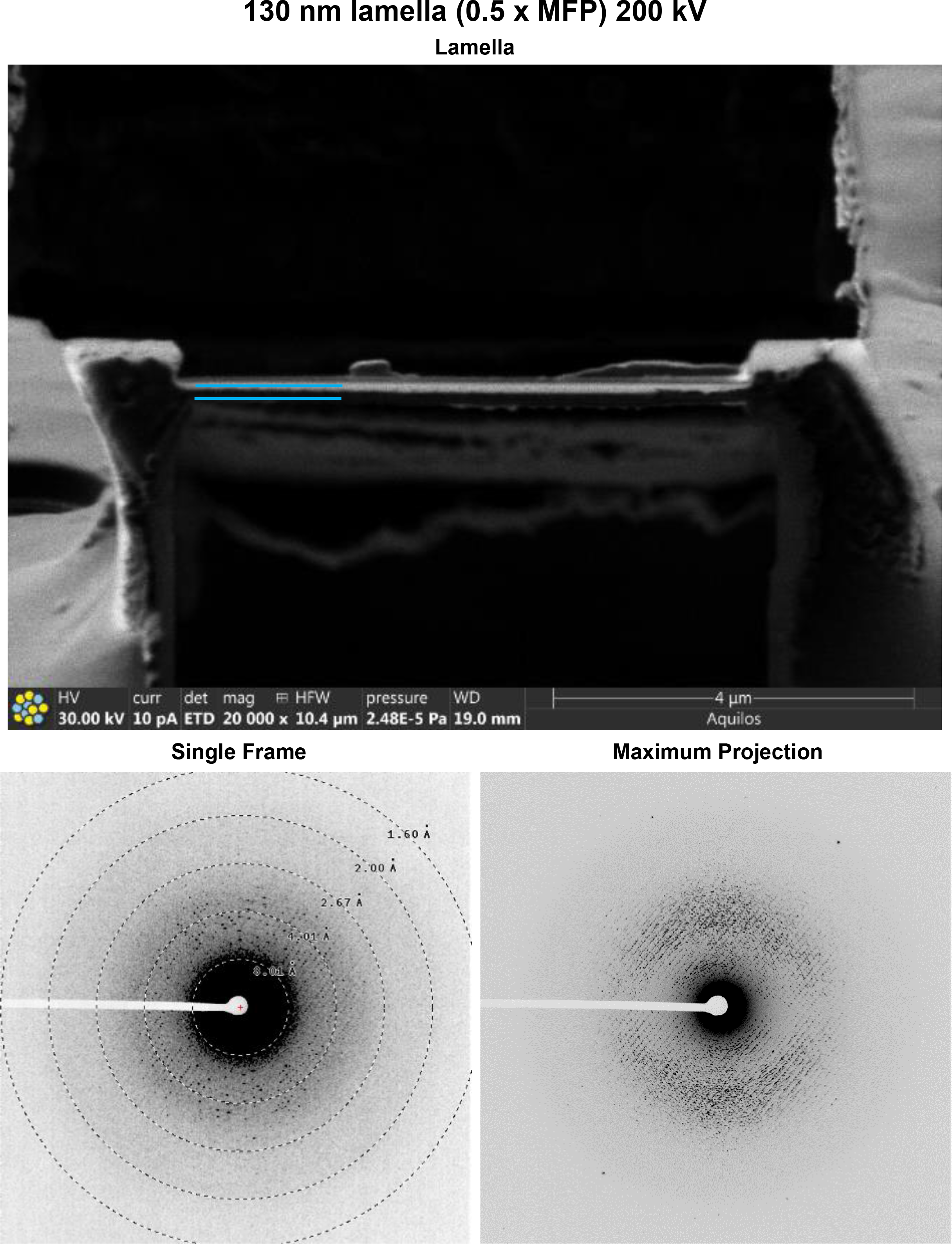
Data from a 130 nm lamella of proteinase K collected at 200 kV. (Top) Image in the FIB directly after the milling finished. (Bottom Left) Single frame of MicroED data (Bottom Right) Maximum projection of the entire MicroED dataset onto a single frame.

**Supplementary Figure 14.**
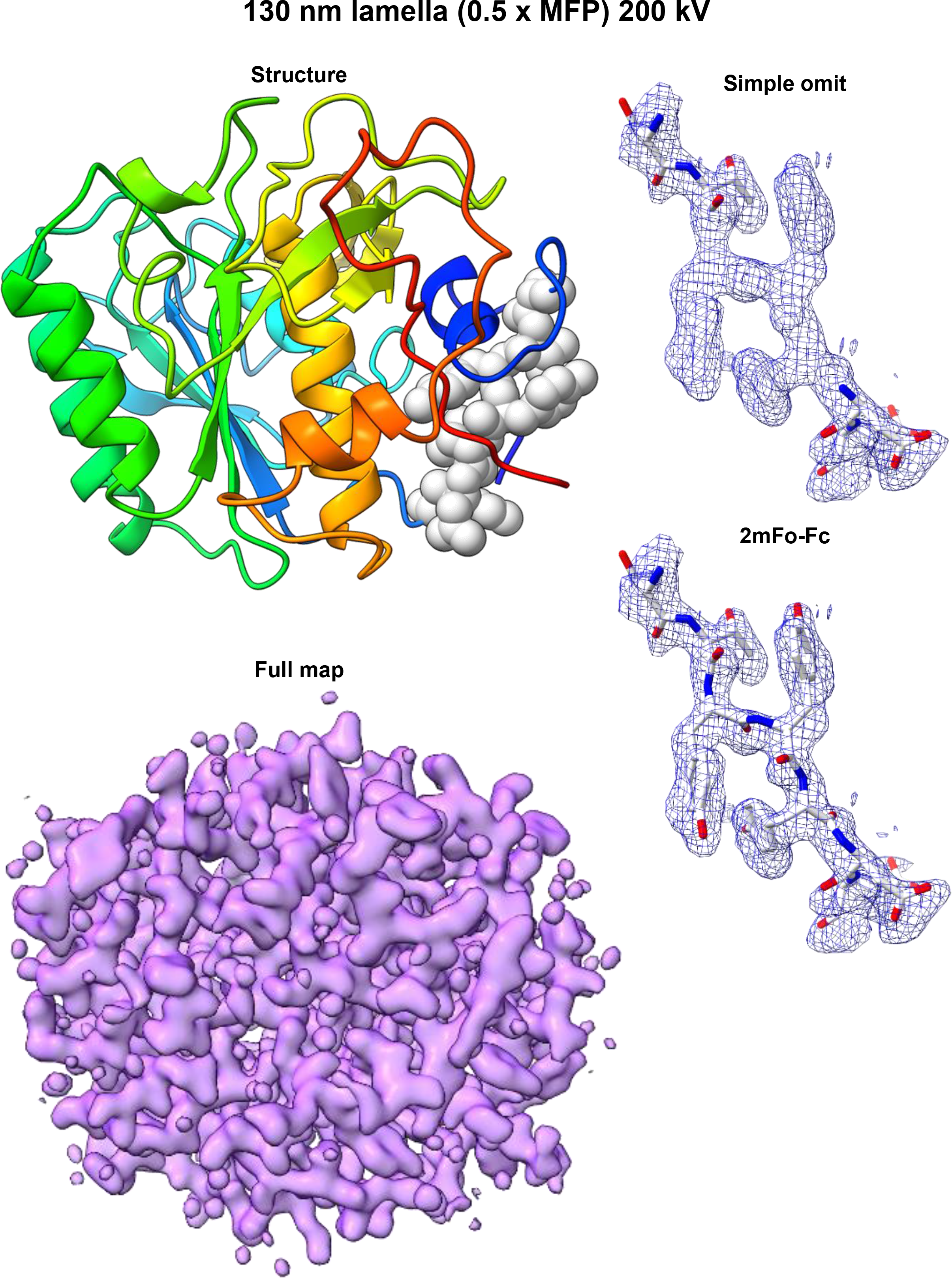
Structure of Proteinase K determined from a 130 nm lamella at 200 kV. (Left Top) Cartoon structure. (Left Bottom) 2mF_o_-F_c_ map contoured at 1.5σ level. (Right Top) 2mF_o_-F_c_ map of the structure generated without three loop residues indicated in the structure on the right. (Right bottom) 2mF_o_-F_c_ map for the same region calculated with the missing residues.

**Supplementary Figure 15.**
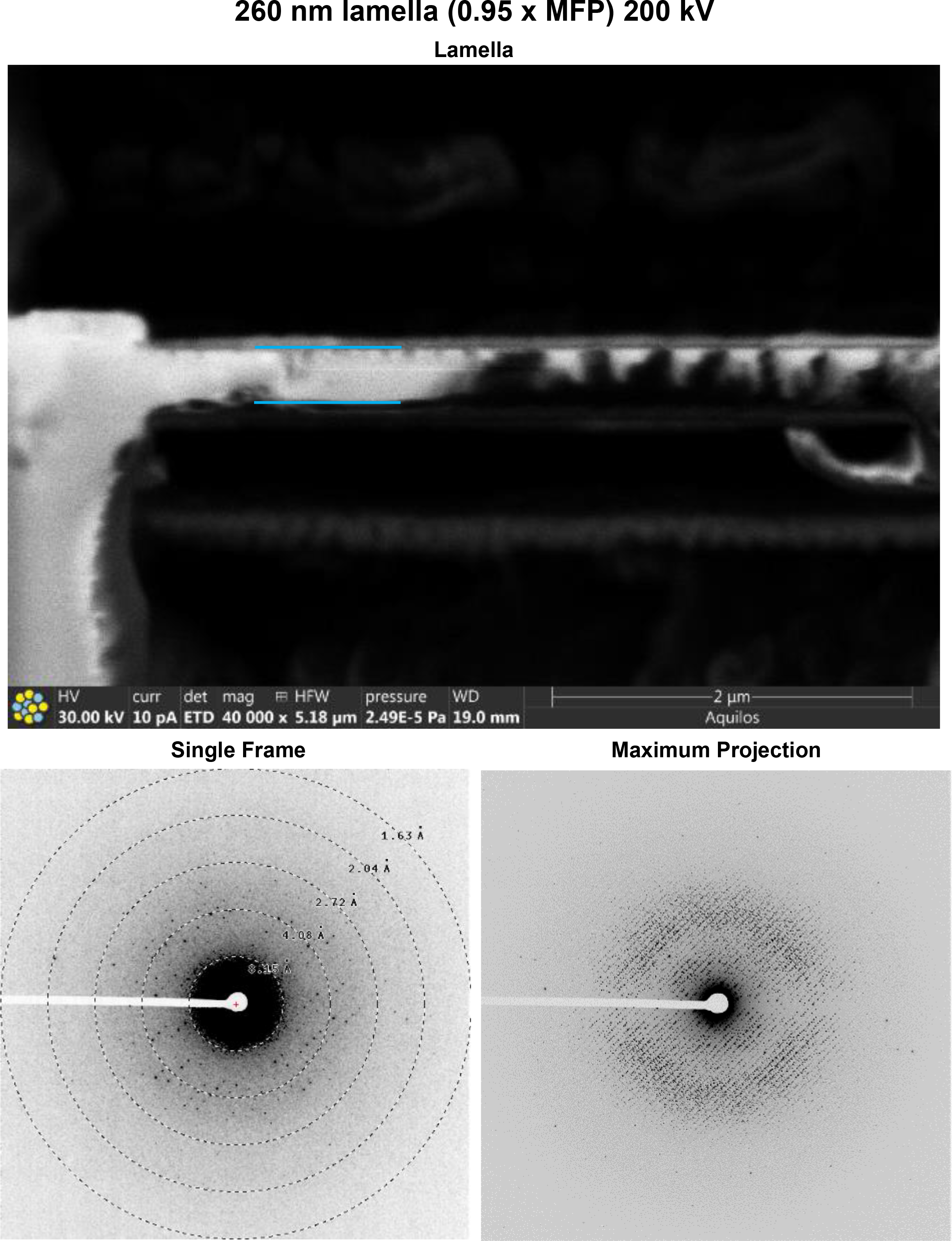
Data from a 260 nm lamella of proteinase K collected at 200 kV. (Top) Image in the FIB directly after the milling finished. (Bottom Left) Single frame of MicroED data (Bottom Right) Maximum projection of the entire MicroED dataset onto a single frame.

**Supplementary Figure 16.**
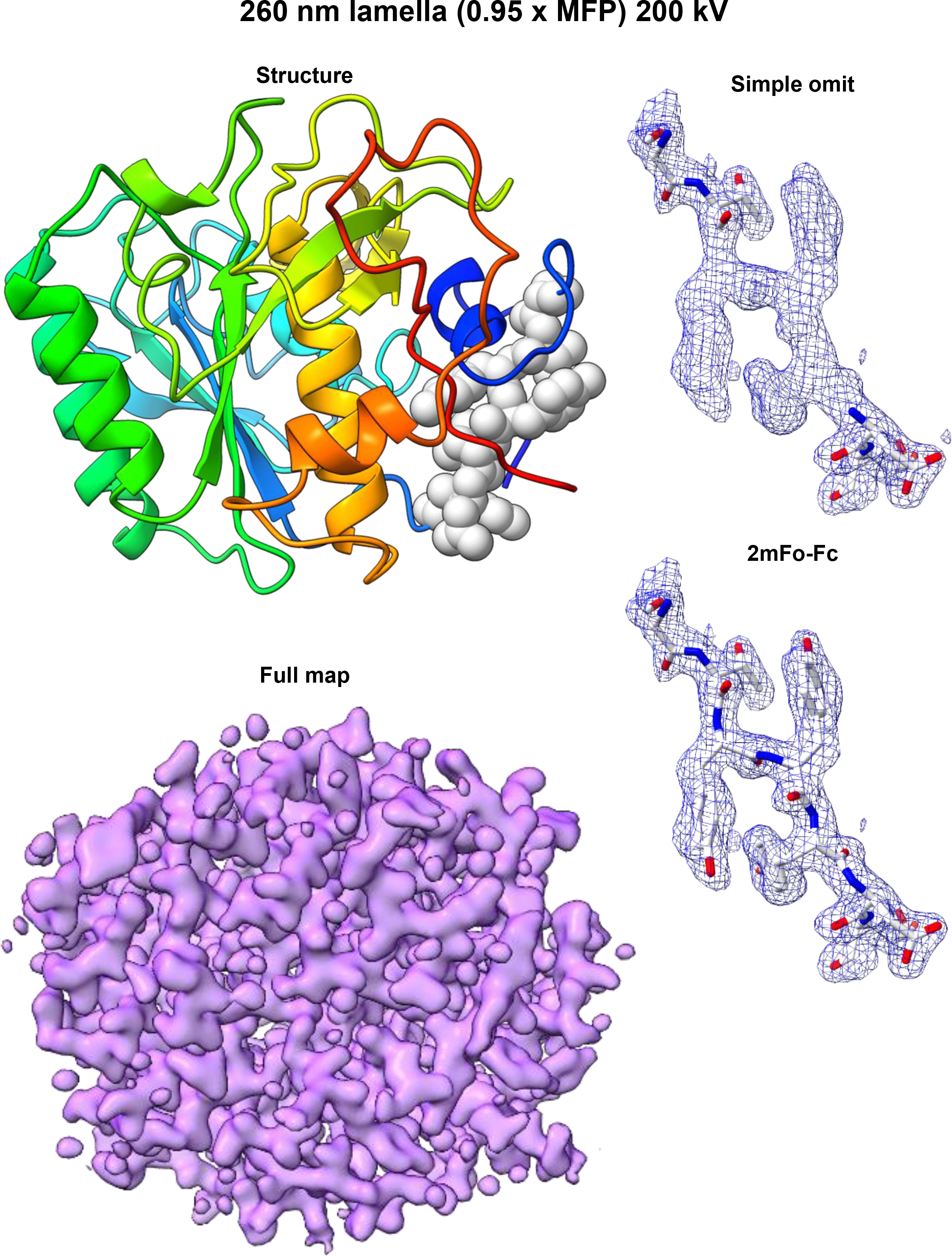
Structure of Proteinase K determined from a 260 nm lamella at 200 kV. (Left Top) Cartoon structure. (Left Bottom) 2mF_o_-F_c_ map contoured at 1.5σ level. (Right Top) 2mF_o_-F_c_ map of the structure generated without three loop residues indicated in the structure on the right. (Right bottom) 2mF_o_-F_c_ map for the same region calculated with the missing residues.

**Supplementary Figure 17.**
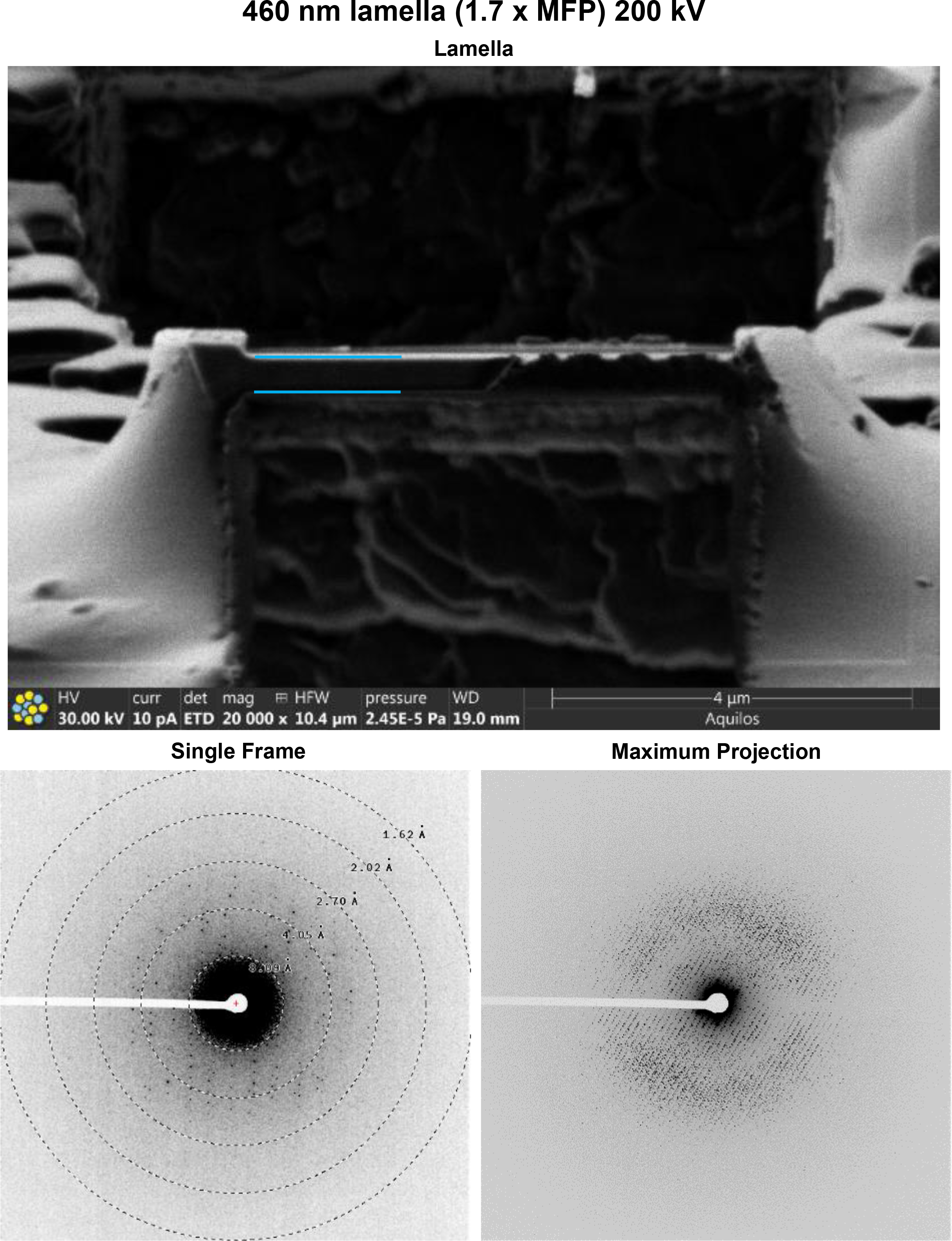
Data from a 460 nm lamella of proteinase K collected at 200 kV. (Top) Image in the FIB directly after the milling finished. (Bottom Left) Single frame of MicroED data (Bottom Right) Maximum projection of the entire MicroED dataset onto a single frame.

**Supplementary Figure 18.**
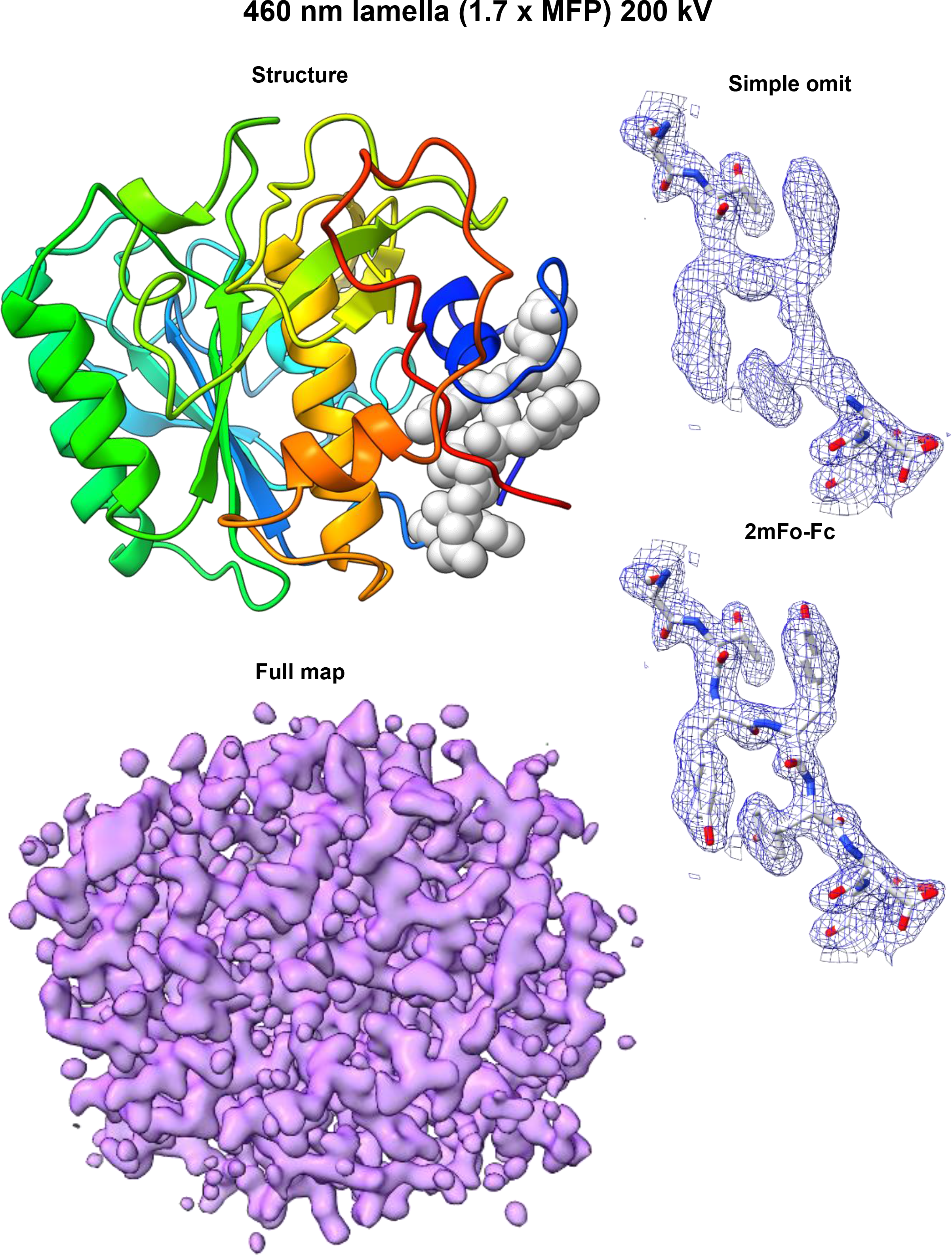
Structure of Proteinase K determined from a 460 nm lamella at 200 kV. (Left Top) Cartoon structure. (Left Bottom) 2mF_o_-F_c_ map contoured at 1.5σ level. (Right Top) 2mF_o_-F_c_ map of the structure generated without three loop residues indicated in the structure on the right. (Right bottom) 2mF_o_-F_c_ map for the same region calculated with the missing residues.

**Supplementary Figure 19.**
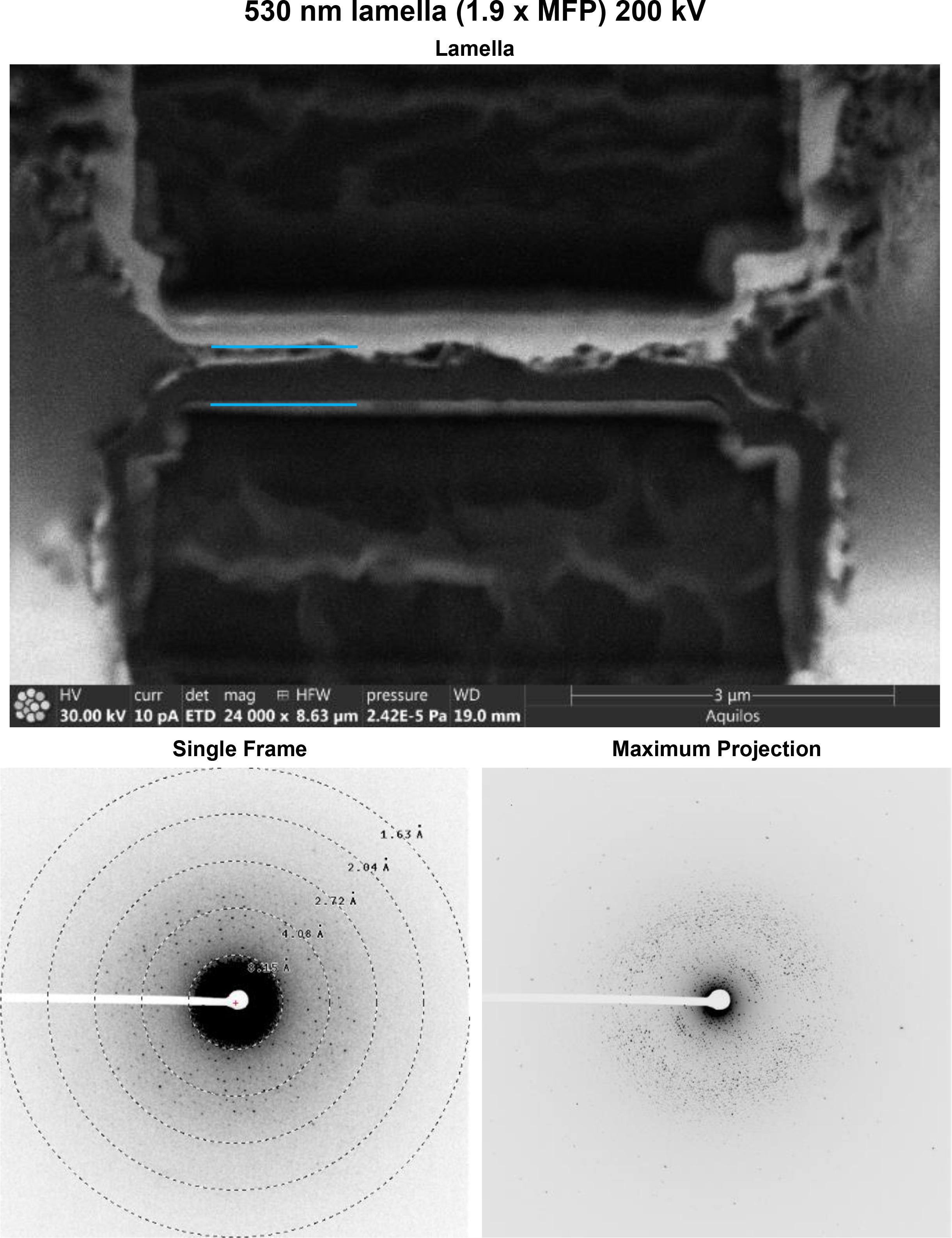
Data from a 530 nm lamella of proteinase K collected at 200 kV. (Top) Image in the FIB directly after the milling finished. (Bottom Left) Single frame of MicroED data (Bottom Right) Maximum projection of the entire MicroED dataset onto a single frame.

**Supplementary Figure 20.**
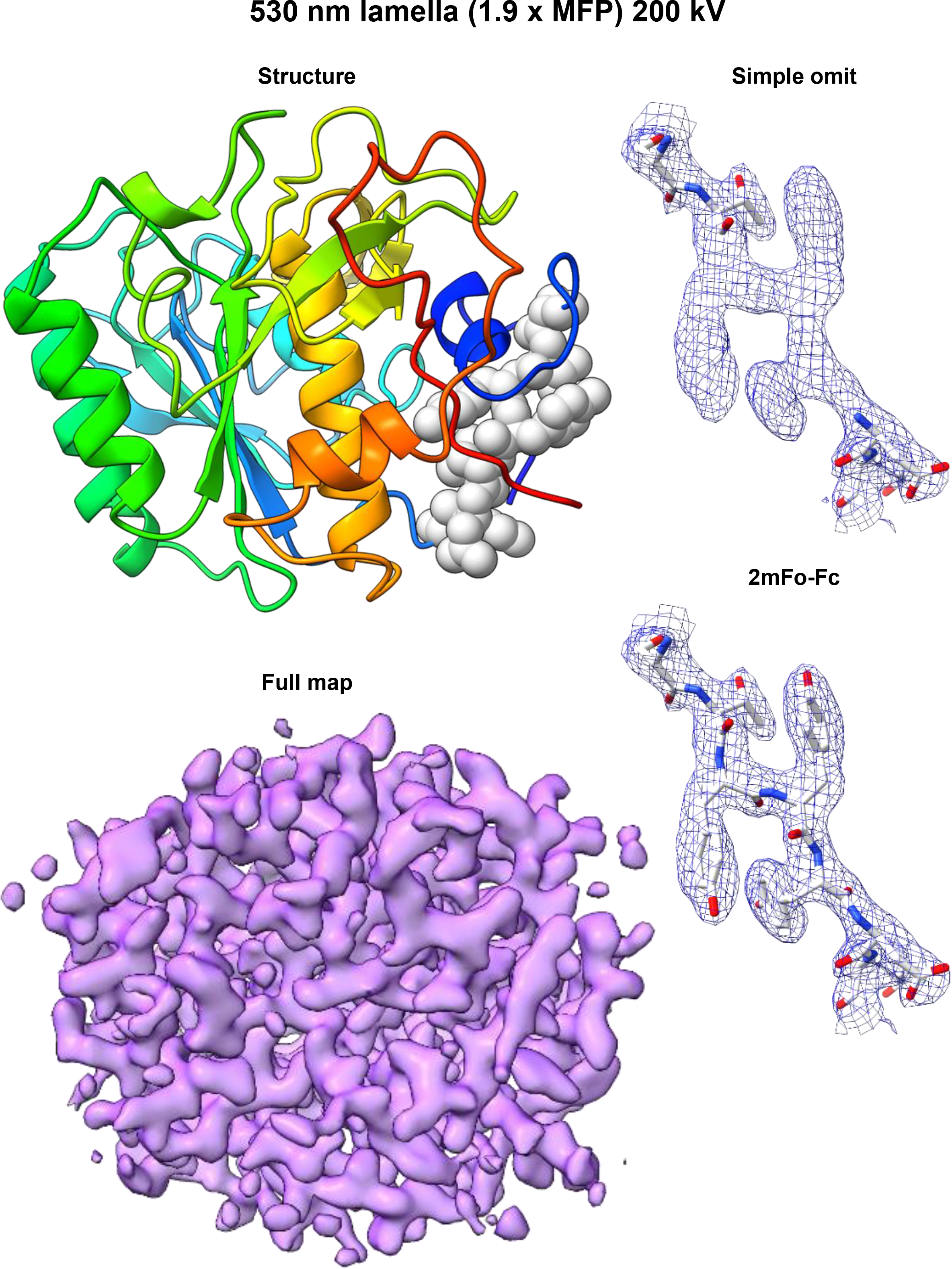
Structure of Proteinase K determined from a 530 nm lamella at 200 kV. (Left Top) Cartoon structure. (Left Bottom) 2mF_o_-F_c_ map contoured at 1.5σ level. (Right Top) 2mF_o_-F_c_ map of the structure generated without three loop residues indicated in the structure on the right. (Right bottom) 2mF_o_-F_c_ map for the same region calculated with the missing residues.

**Supplementary Figure 21.**
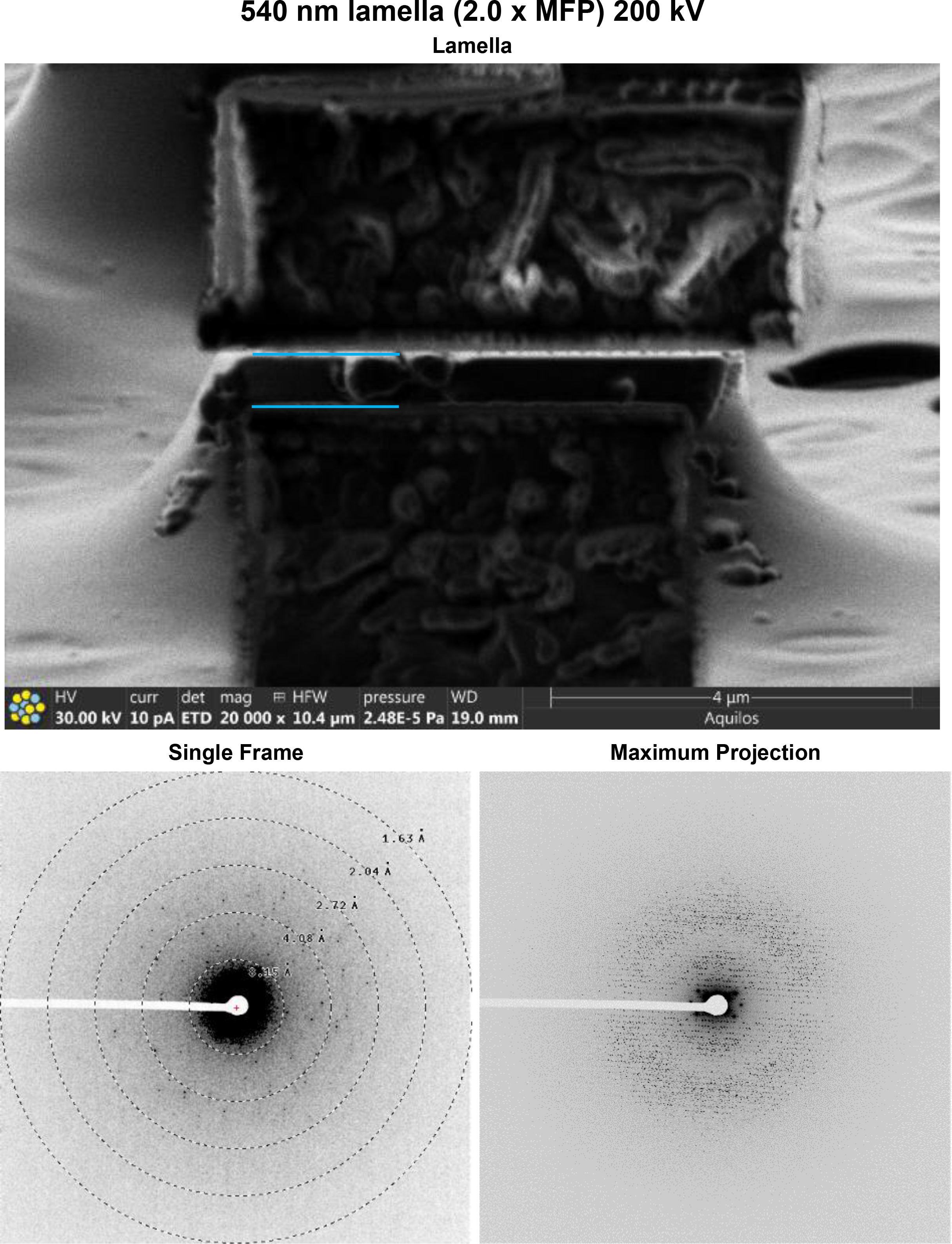
Data from a 540 nm lamella of proteinase K collected at 200 kV. (Top) Image in the FIB directly after the milling finished. (Bottom Left) Single frame of MicroED data (Bottom Right) Maximum projection of the entire MicroED dataset onto a single frame.

**Supplementary Figure 22.**
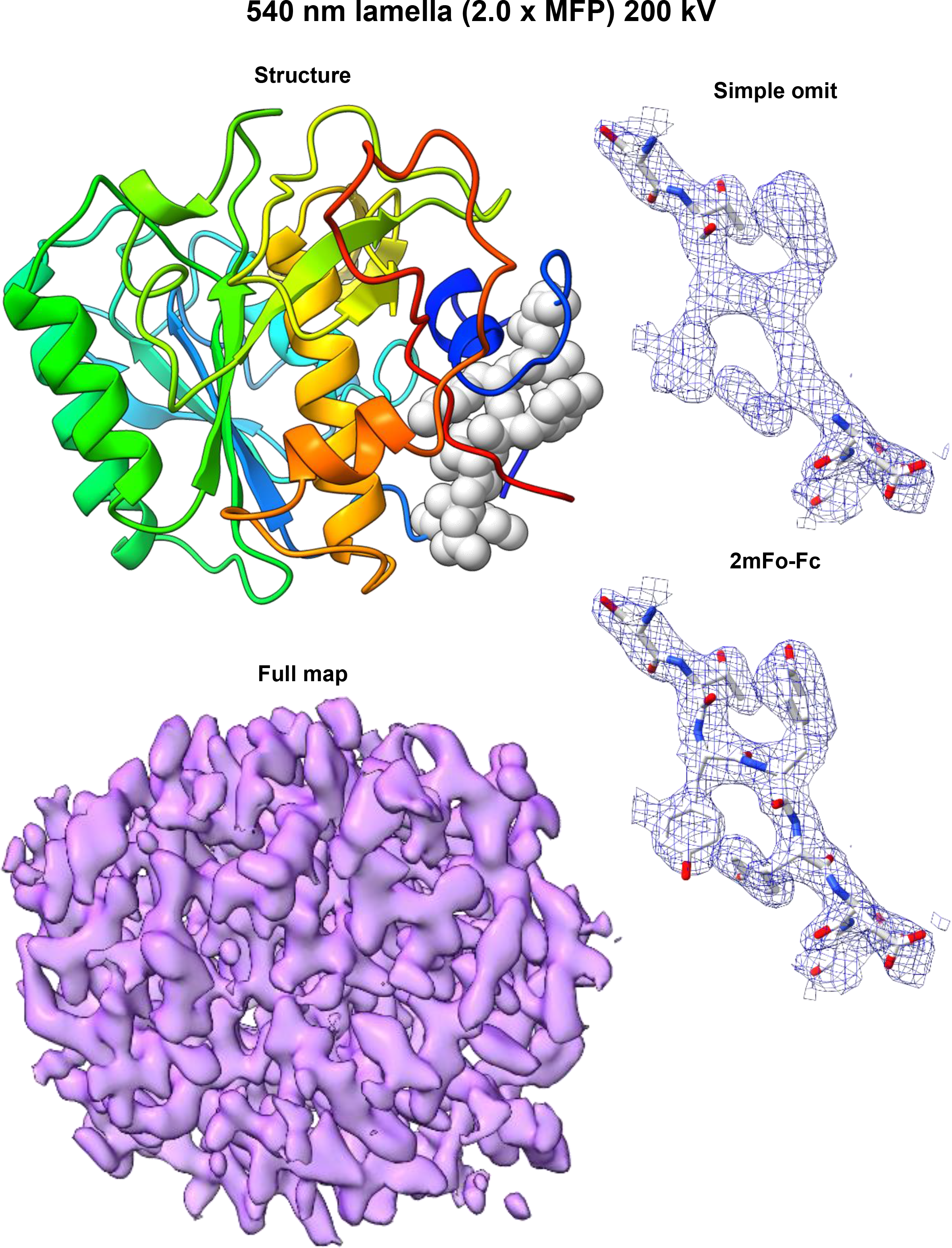
Structure of Proteinase K determined from a 540 nm lamella at 200 kV. (Left Top) Cartoon structure. (Left Bottom) 2mF_o_-F_c_ map contoured at 1.5σ level. (Right Top) 2mF_o_-F_c_ map of the structure generated without three loop residues indicated in the structure on the right. (Right bottom) 2mF_o_-F_c_ map for the same region calculated with the missing residues.

**Supplementary Figure 23.**
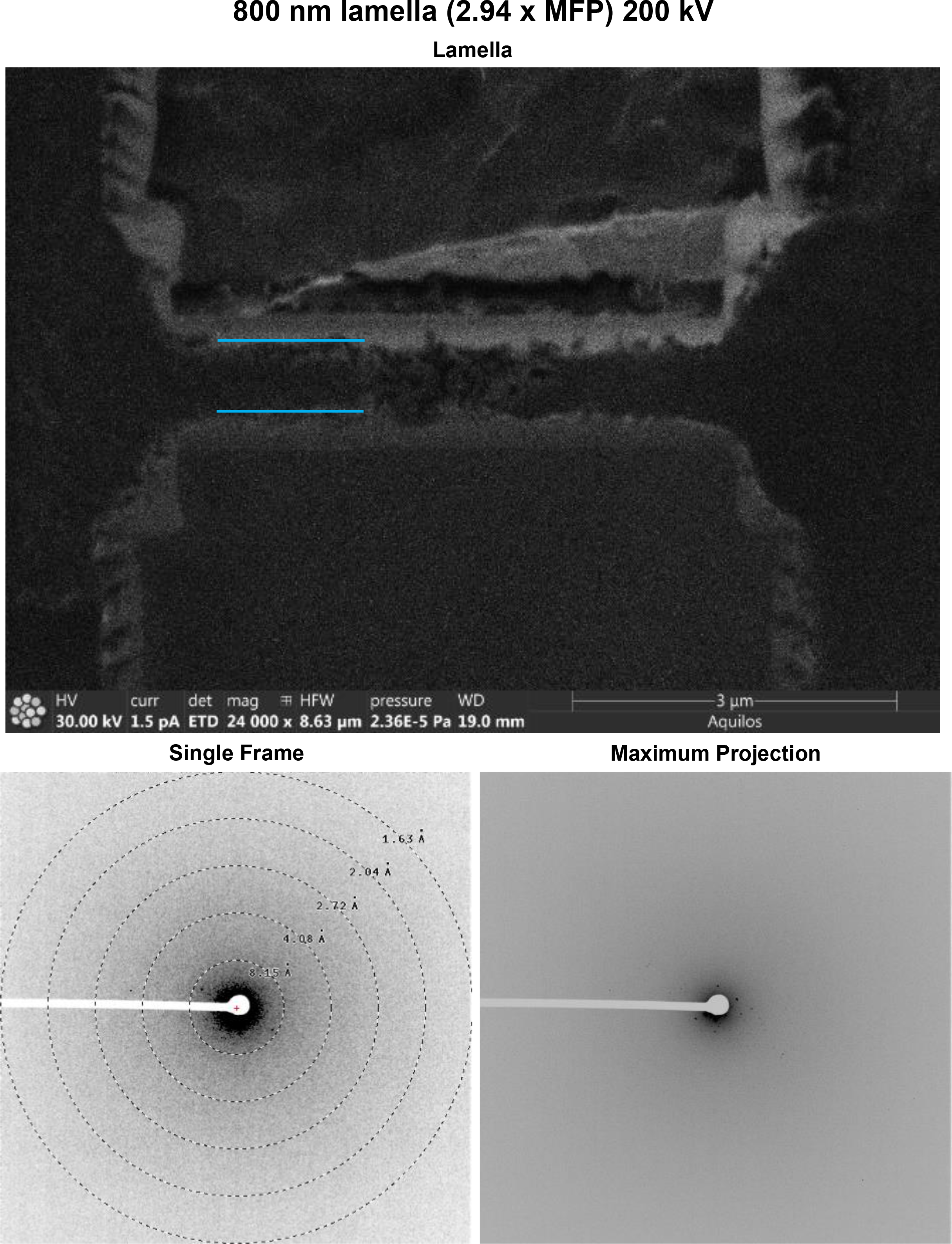
Data from an 800 nm lamella of proteinase K collected at 200 kV. (Top) Image in the FIB directly after the milling finished. (Bottom Left) Single frame of MicroED data (Bottom Right) Maximum projection of the entire MicroED dataset onto a single frame.

**Supplementary Figure 24.**
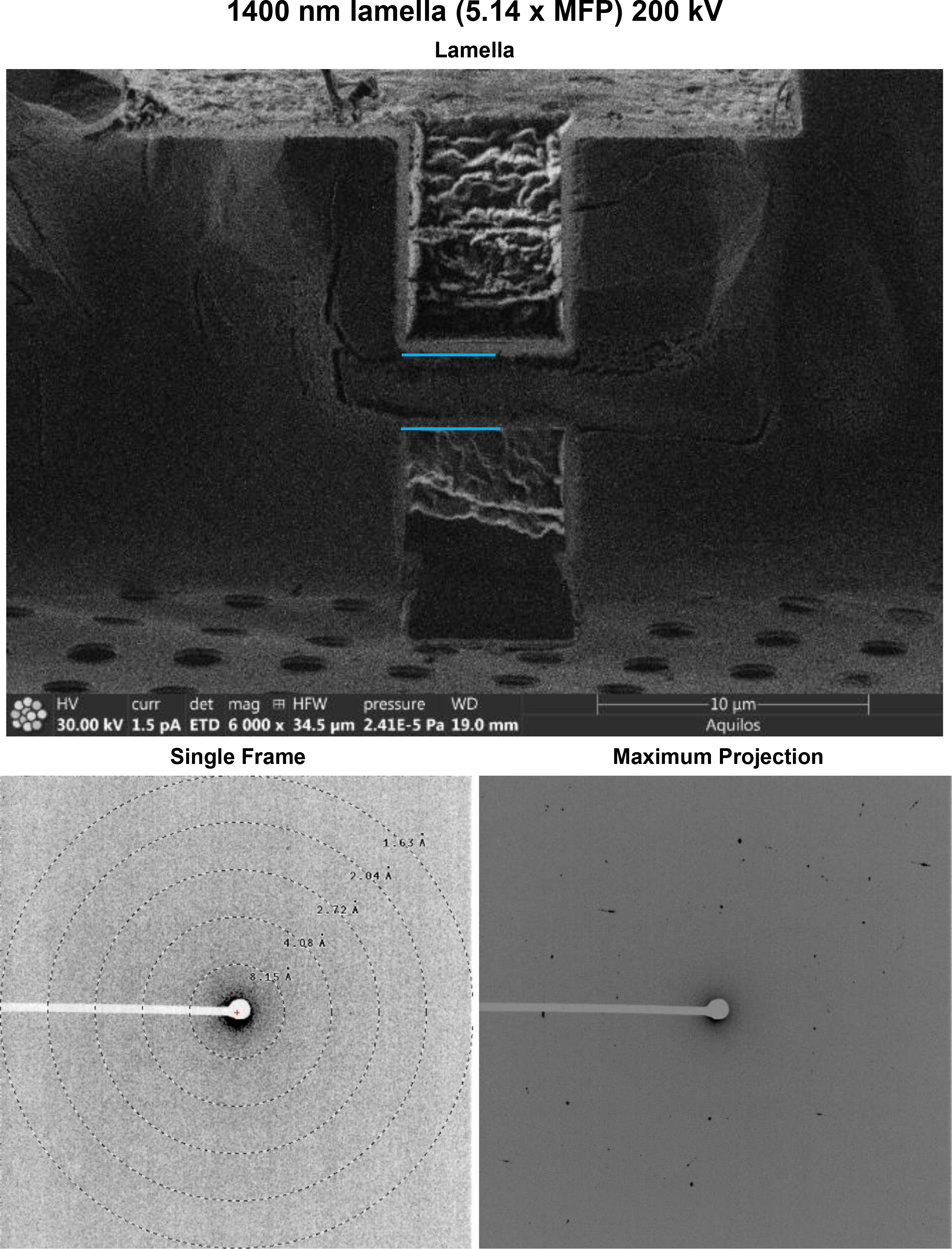
Data from a 1400 nm lamella of proteinase K collected at 200 kV. (Top) Image in the FIB directly after the milling finished. (Bottom Left) Single frame of MicroED data (Bottom Right) Maximum projection of the entire MicroED dataset onto a single frame.

**Supplementary Figure 25.**
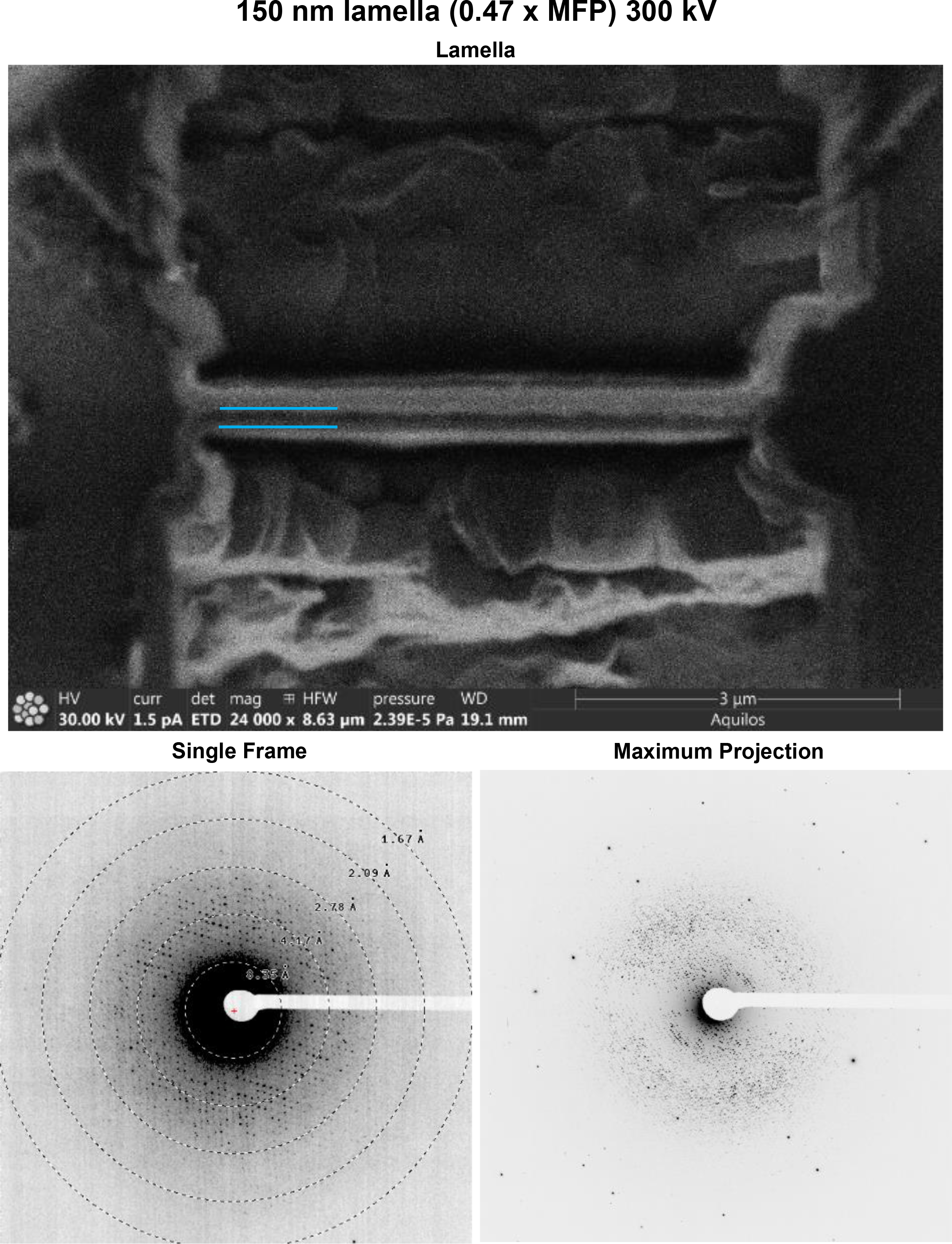
Data from a 150 nm lamella of proteinase K collected at 300 kV. (Top) Image in the FIB directly after the milling finished. (Bottom Left) Single frame of MicroED data (Bottom Right) Maximum projection of the entire MicroED dataset onto a single frame.

**Supplementary Figure 26.**
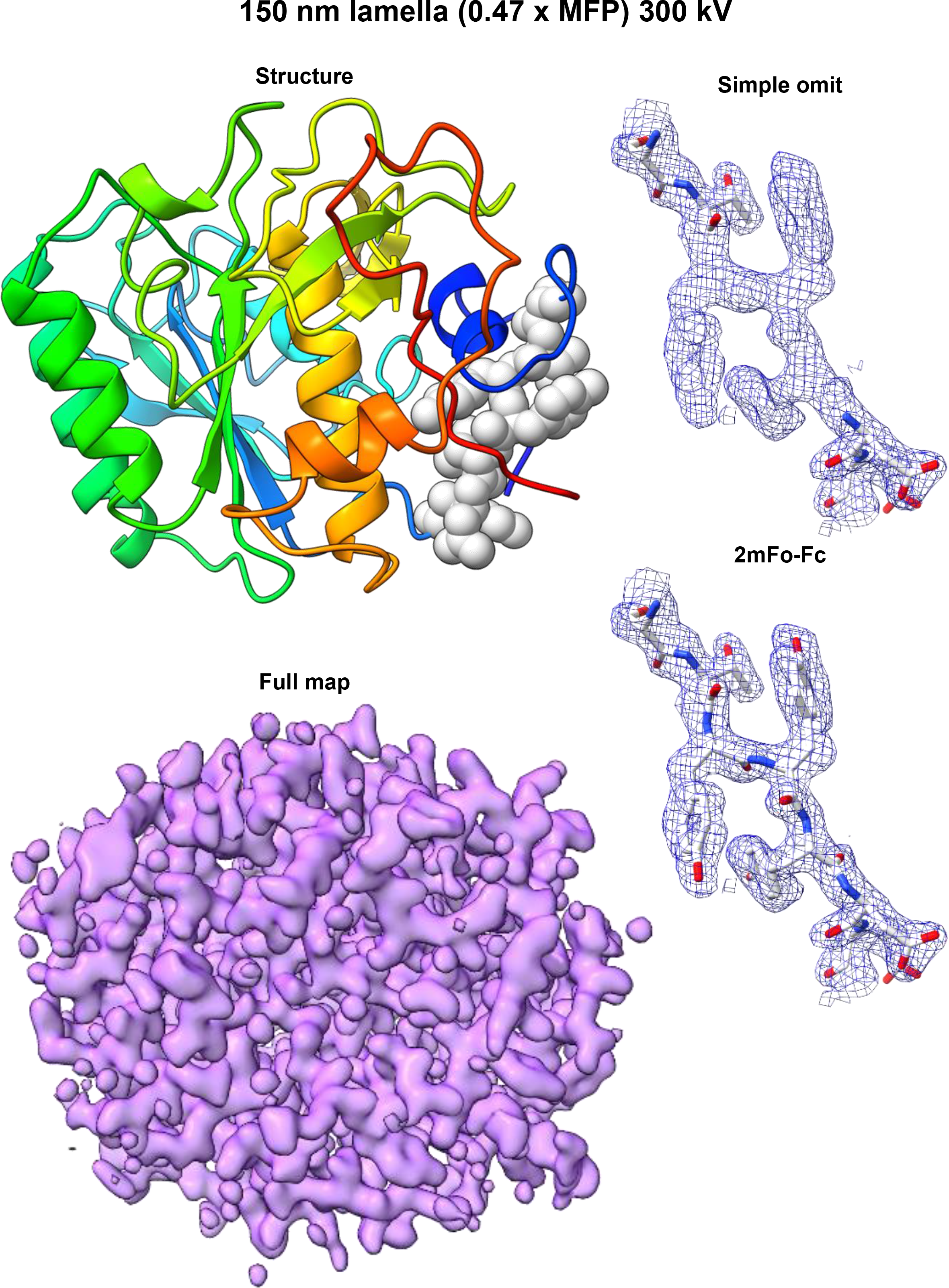
Structure of Proteinase K determined from a 150 nm lamella at 300 kV. (Left Top) Cartoon structure. (Left Bottom) 2mF_o_-F_c_ map contoured at 1.5σ level. (Right Top) 2mF_o_-F_c_ map of the structure generated without three loop residues indicated in the structure on the right. (Right bottom) 2mF_o_-F_c_ map for the same region calculated with the missing residues.

**Supplementary Figure 27.**
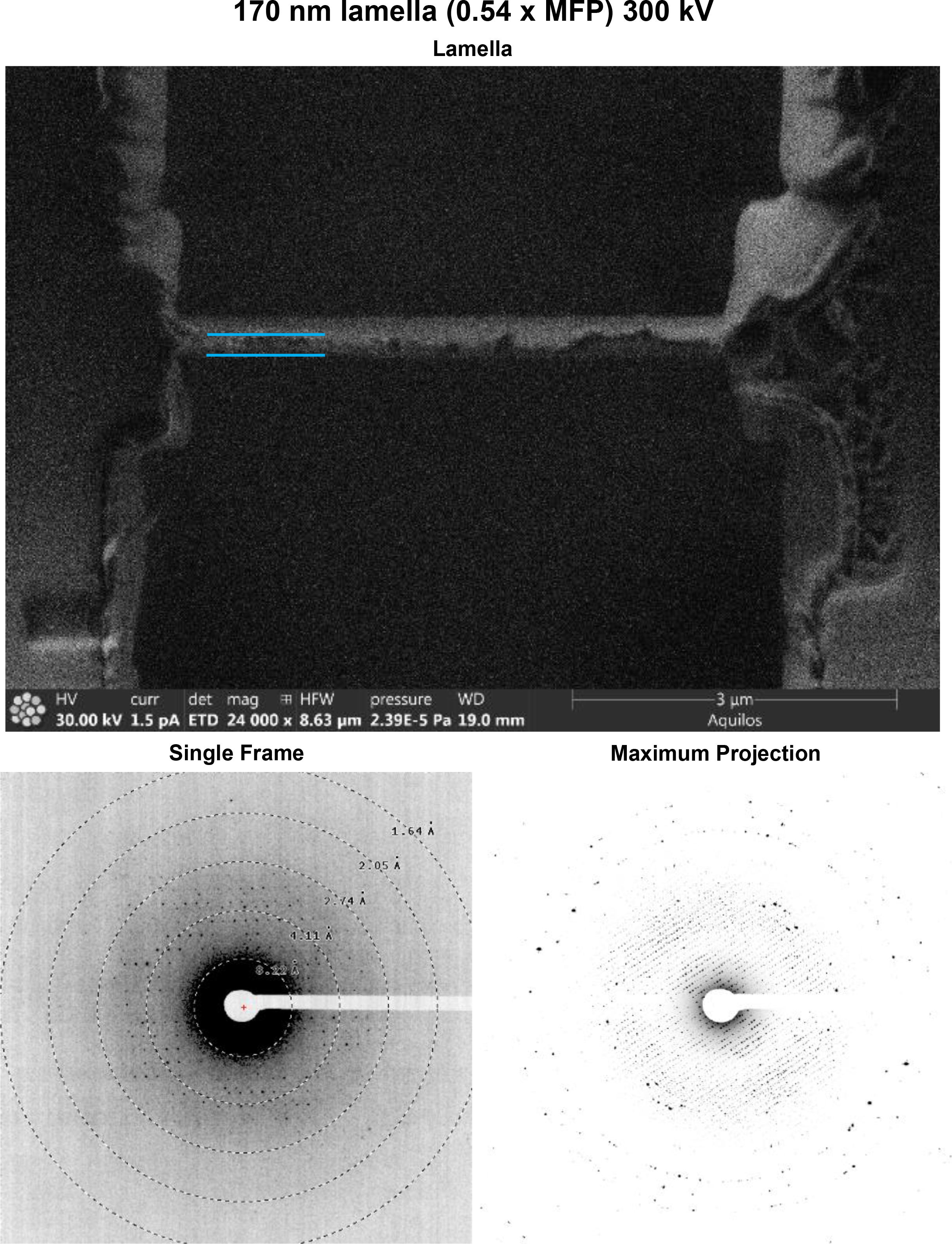
Data from a 170 nm lamella of proteinase K collected at 300 kV. (Top) Image in the FIB directly after the milling finished. (Bottom Left) Single frame of MicroED data (Bottom Right) Maximum projection of the entire MicroED dataset onto a single frame.

**Supplementary Figure 28.**
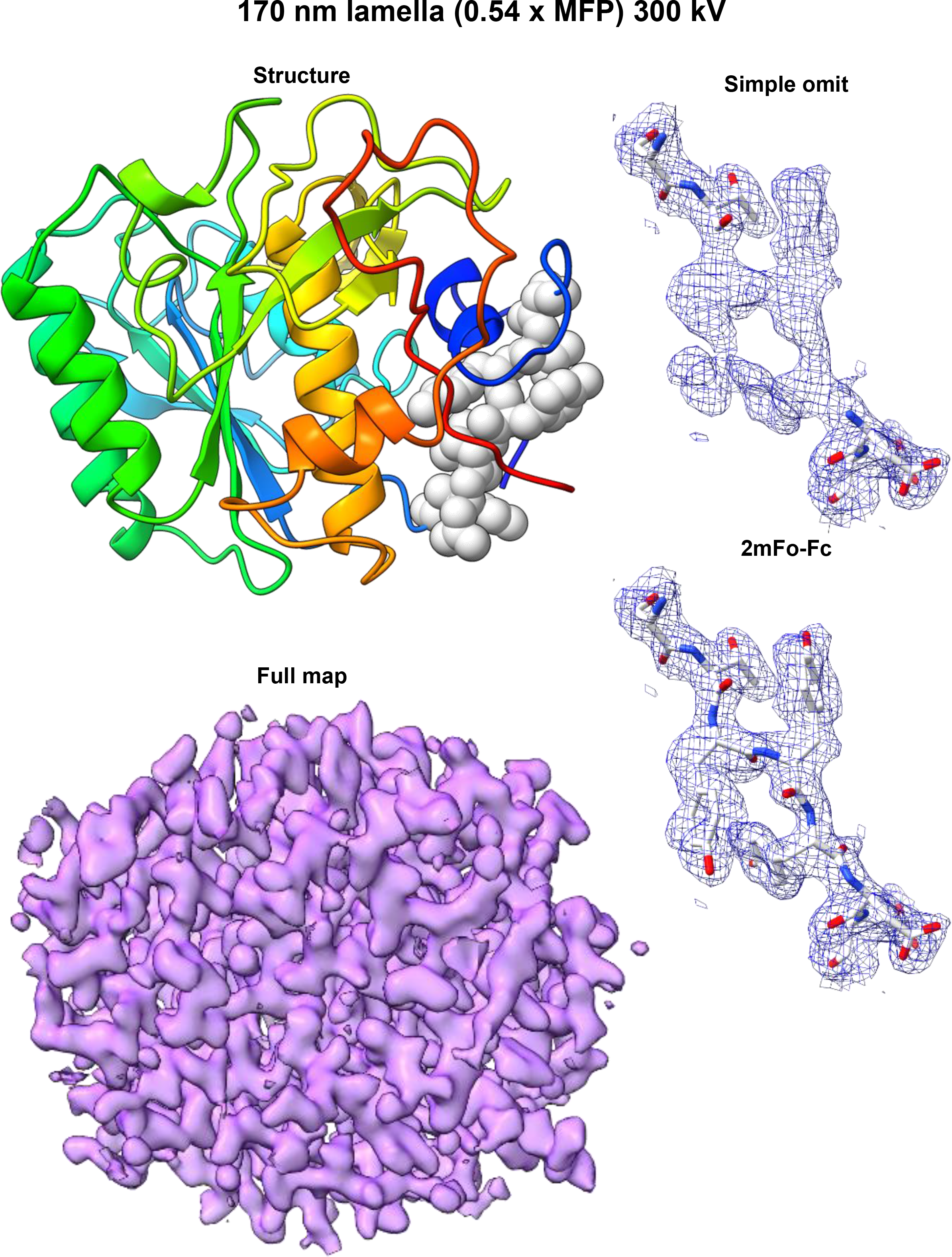
Structure of Proteinase K determined from a 170 nm lamella at 300 kV. (Left Top) Cartoon structure. (Left Bottom) 2mF_o_-F_c_ map contoured at 1.5σ level. (Right Top) 2mF_o_-F_c_ map of the structure generated without three loop residues indicated in the structure on the right. (Right bottom) 2mF_o_-F_c_ map for the same region calculated with the missing residues.

**Supplementary Figure 29.**
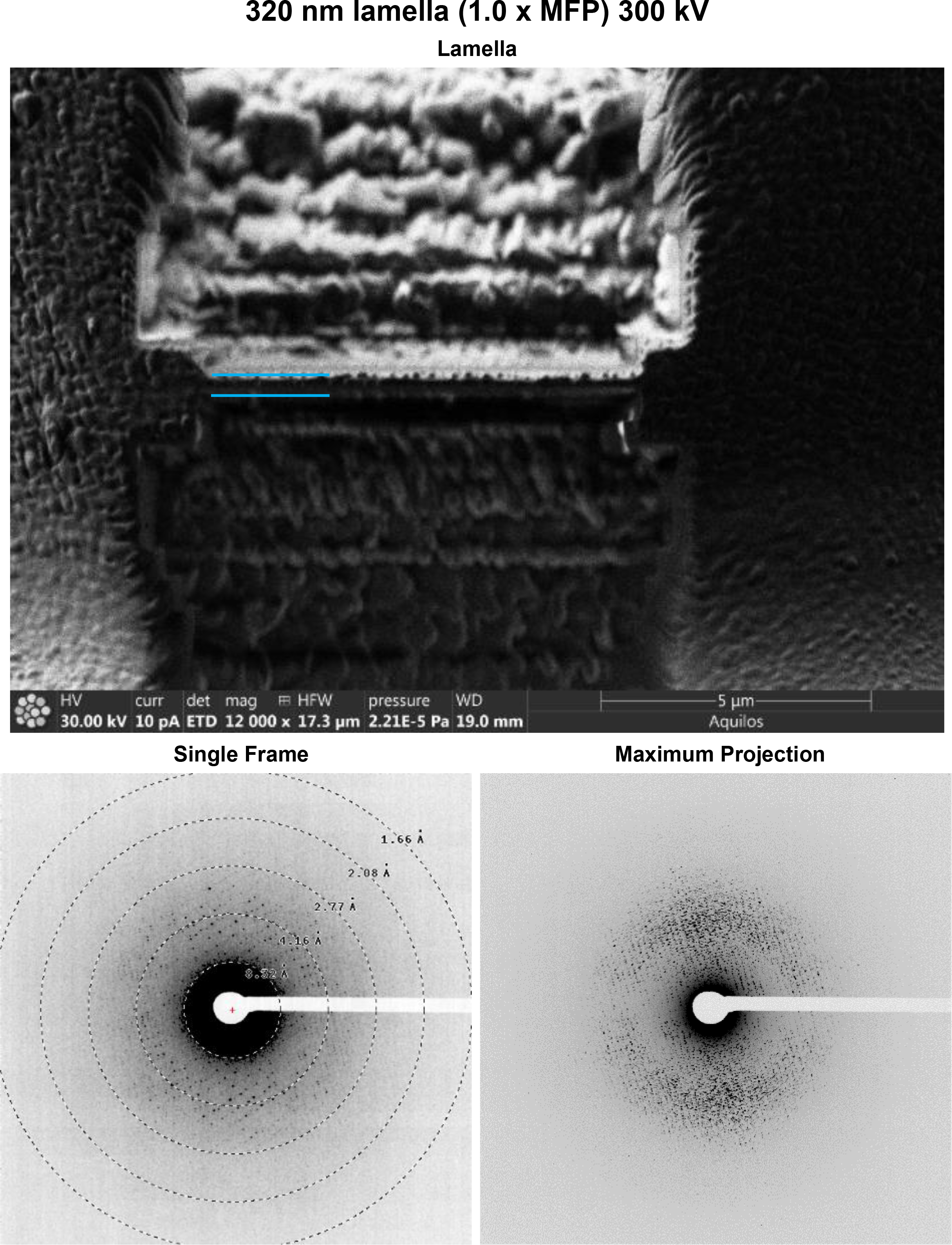
Data from a 320 nm lamella of proteinase K collected at 300 kV. (Top) Image in the FIB directly after the milling finished. (Bottom Left) Single frame of MicroED data (Bottom Right) Maximum projection of the entire MicroED dataset onto a single frame.

**Supplementary Figure 30.**
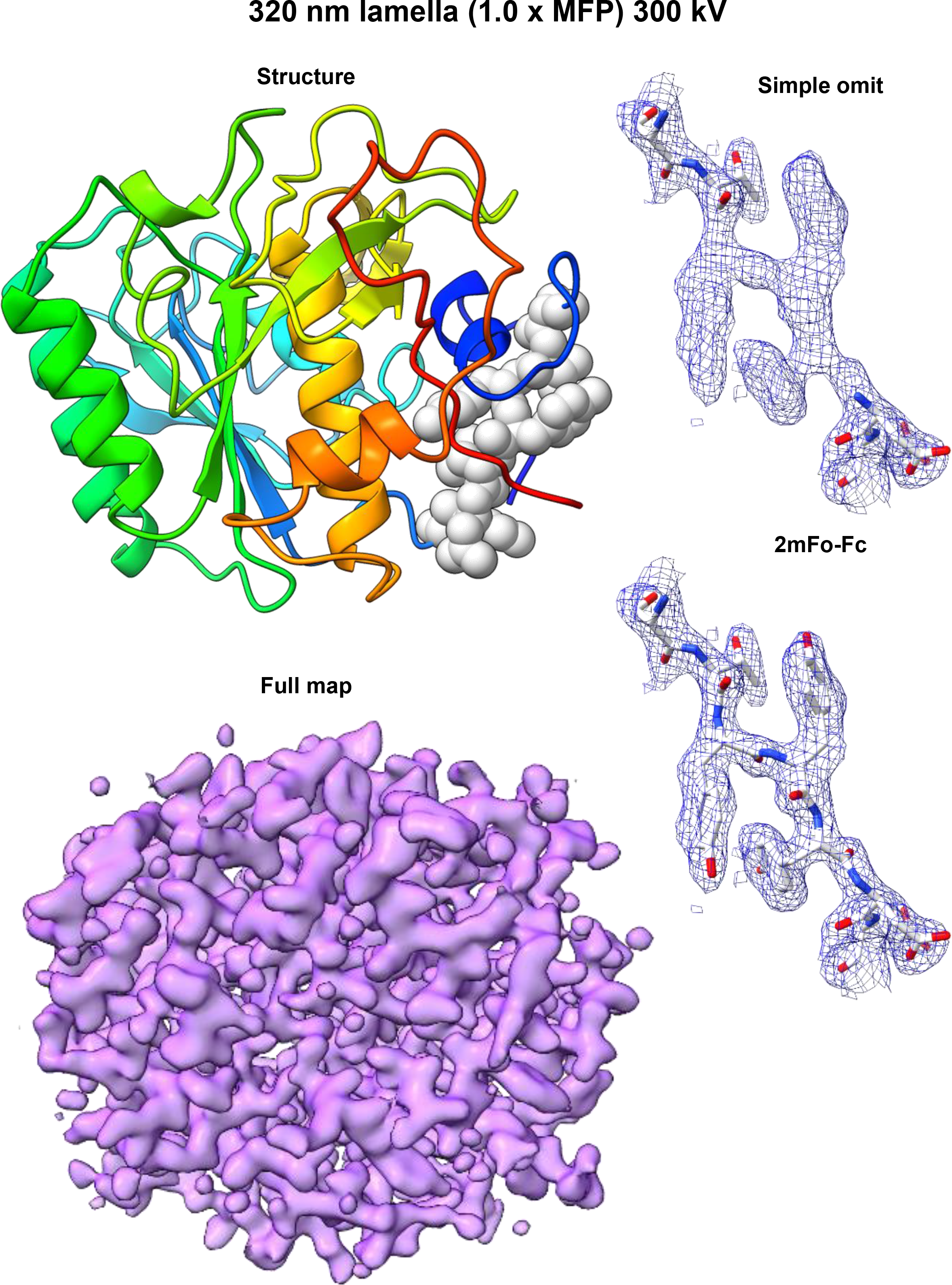
Structure of Proteinase K determined from a 320 nm lamella at 300 kV. (Left Top) Cartoon structure. (Left Bottom) 2mF_o_-F_c_ map contoured at 1.5σ level. (Right Top) 2mF_o_-F_c_ map of the structure generated without three loop residues indicated in the structure on the right. (Right bottom) 2mF_o_-F_c_ map for the same region calculated with the missing residues.

**Supplementary Figure 31.**
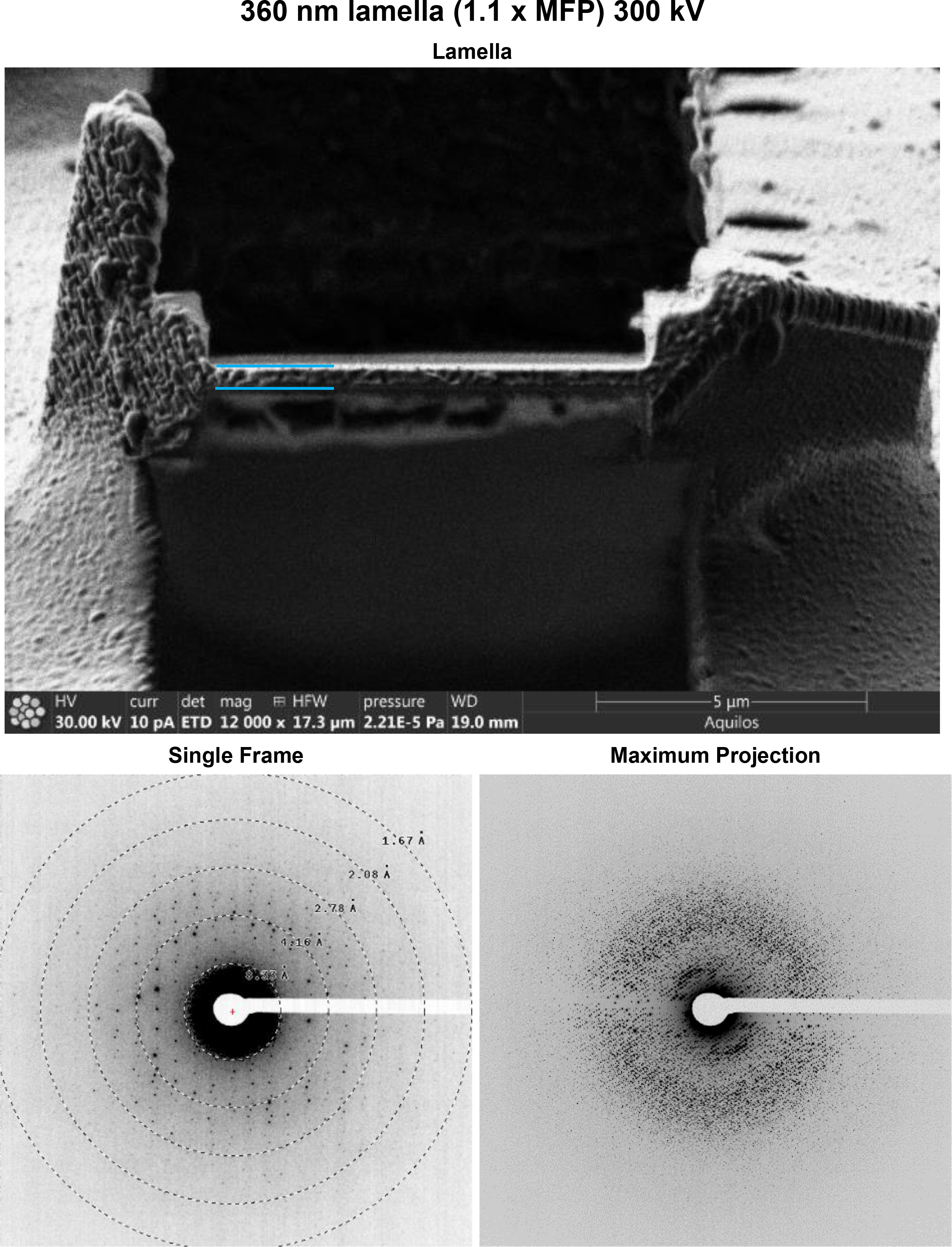
Data from a 360 nm lamella of proteinase K collected at 300 kV. (Top) Image in the FIB directly after the milling finished. (Bottom Left) Single frame of MicroED data (Bottom Right) Maximum projection of the entire MicroED dataset onto a single frame.

**Supplementary Figure 32.**
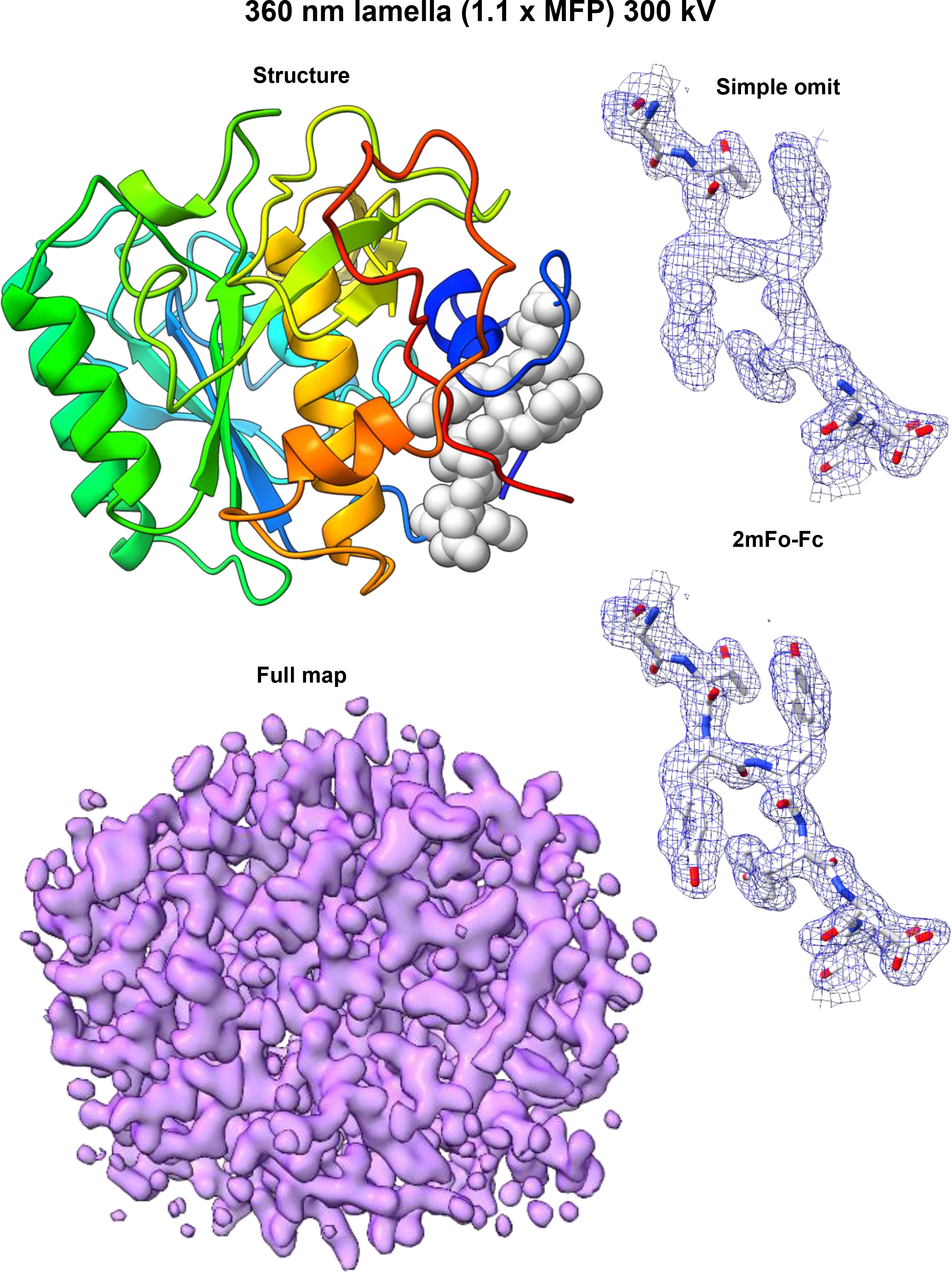
Structure of Proteinase K determined from a 360 nm lamella at 300 kV. (Left Top) Cartoon structure. (Left Bottom) 2mF_o_-F_c_ map contoured at 1.5σ level. (Right Top) 2mF_o_-F_c_ map of the structure generated without three loop residues indicated in the structure on the right. (Right bottom) 2mF_o_-F_c_ map for the same region calculated with the missing residues.

**Supplementary Figure 33.**
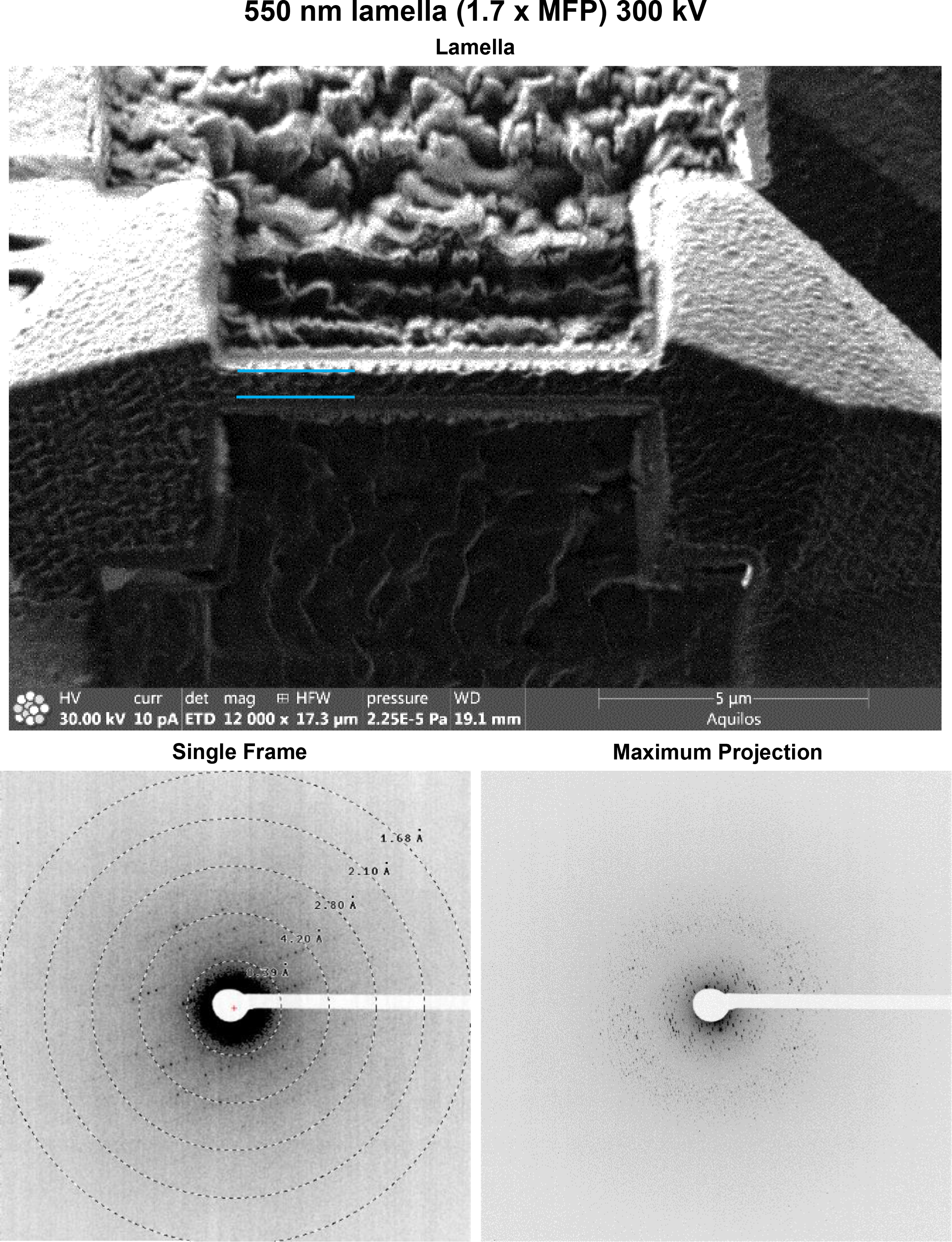
Data from a 550 nm lamella of proteinase K collected at 300 kV. (Top) Image in the FIB directly after the milling finished. (Bottom Left) Single frame of MicroED data (Bottom Right) Maximum projection of the entire MicroED dataset onto a single frame.

**Supplementary Figure 34.**
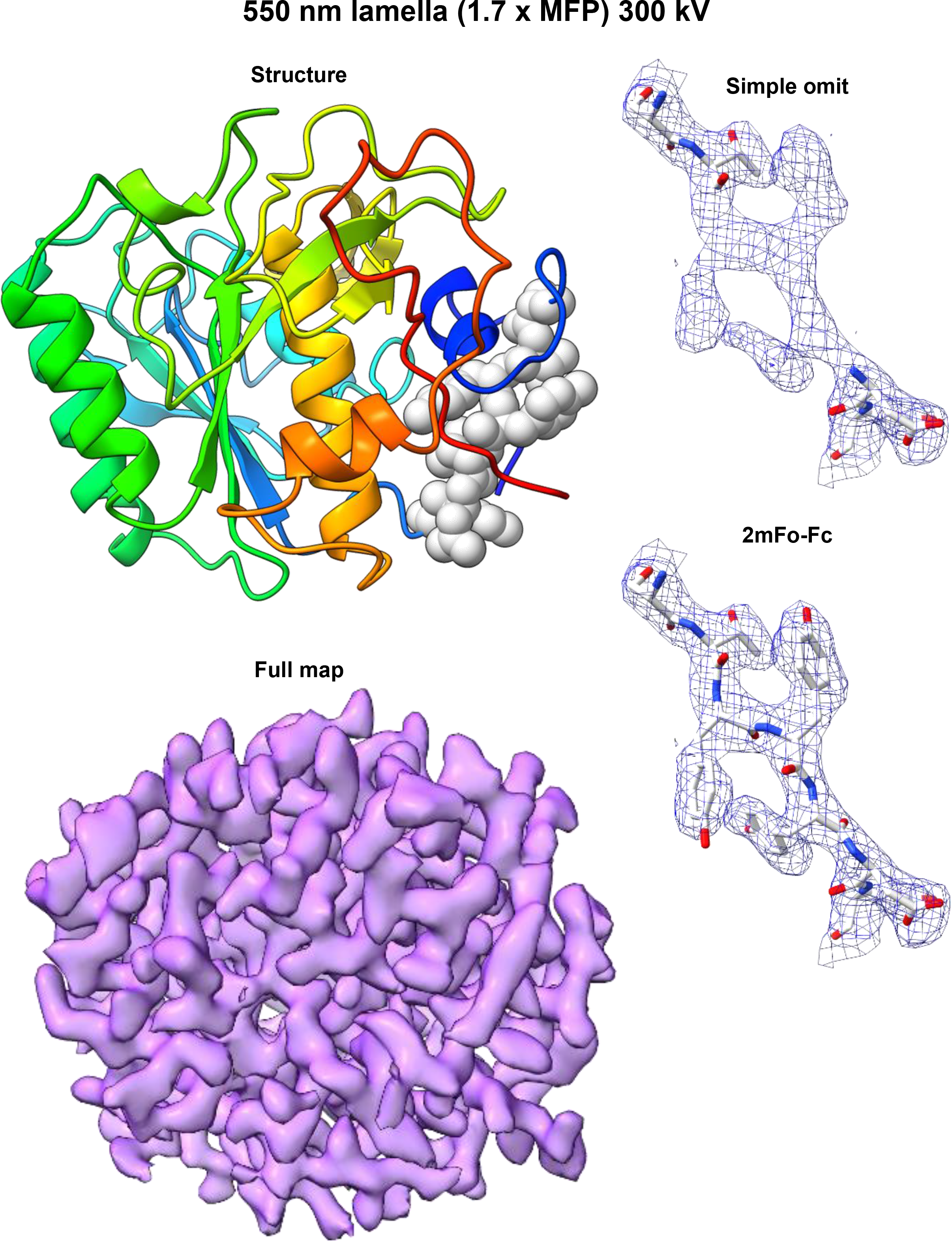
Structure of Proteinase K determined from a 550 nm lamella at 300 kV. (Left Top) Cartoon structure. (Left Bottom) 2mF_o_-F_c_ map contoured at 1.5σ level. (Right Top) 2mF_o_-F_c_ map of the structure generated without three loop residues indicated in the structure on the right. (Right bottom) 2mF_o_-F_c_ map for the same region calculated with the missing residues.

**Supplementary Figure 35.**
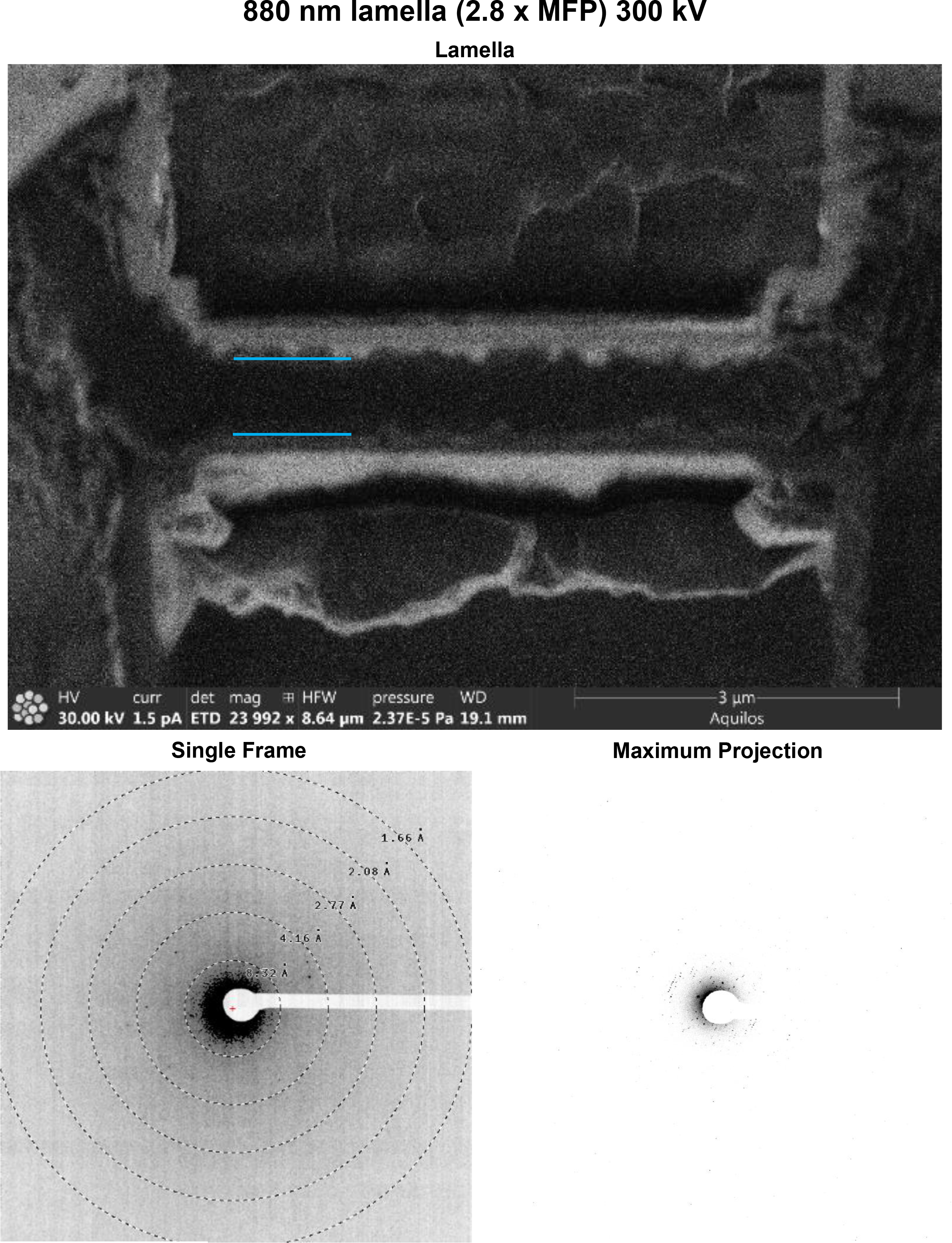
Data from a 880 nm lamella of proteinase K collected at 300 kV. (Top) Image in the FIB directly after the milling finished. (Bottom Left) Single frame of MicroED data (Bottom Right) Maximum projection of the entire MicroED dataset onto a single frame.

**Supplementary Figure 36.**
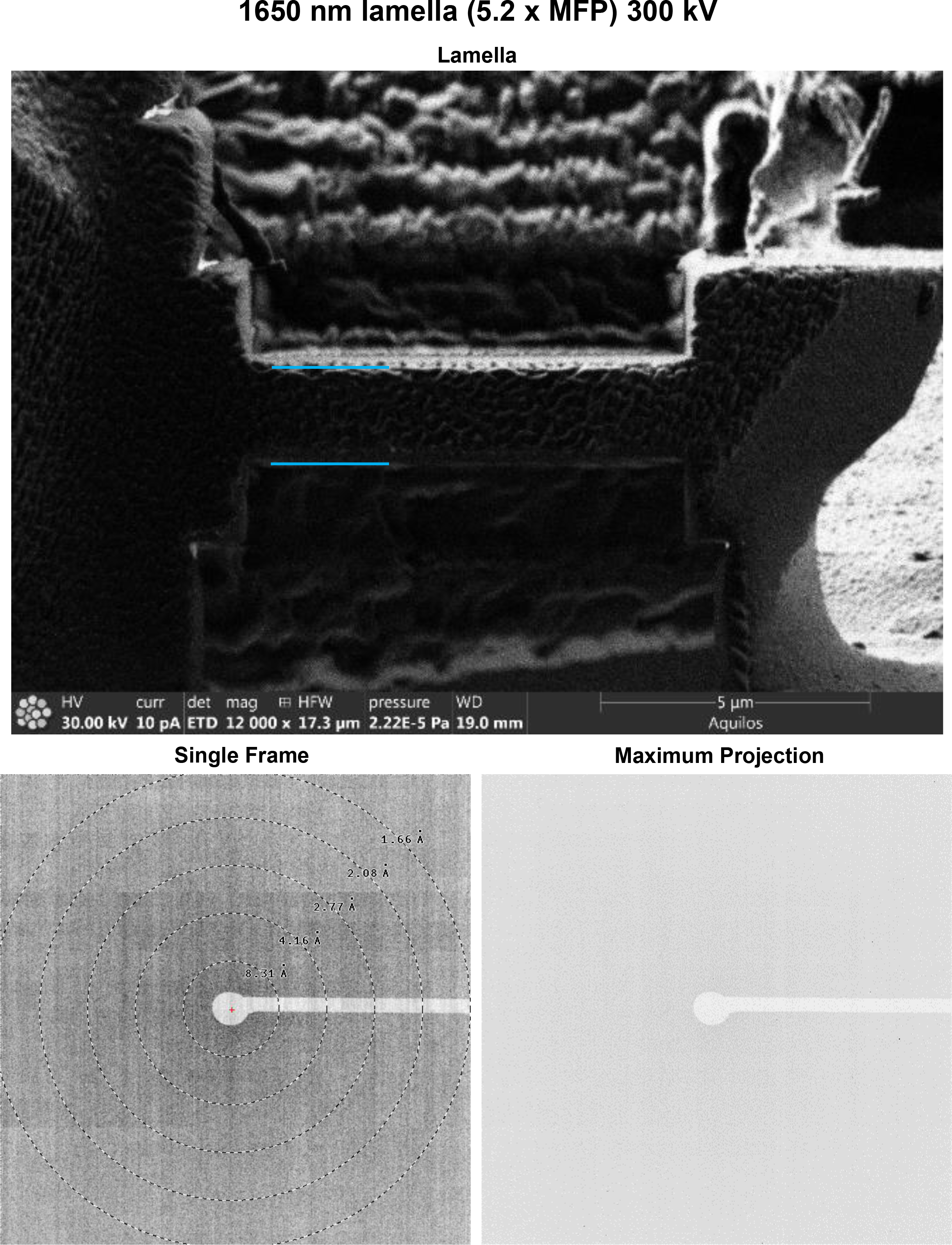
Data from a 1650 nm lamella of proteinase K collected at 300 kV. (Top) Image in the FIB directly after the milling finished. (Bottom Left) Single frame of MicroED data (Bottom Right) Maximum projection of the entire MicroED dataset onto a single frame.

**Supplementary Figure 37.**
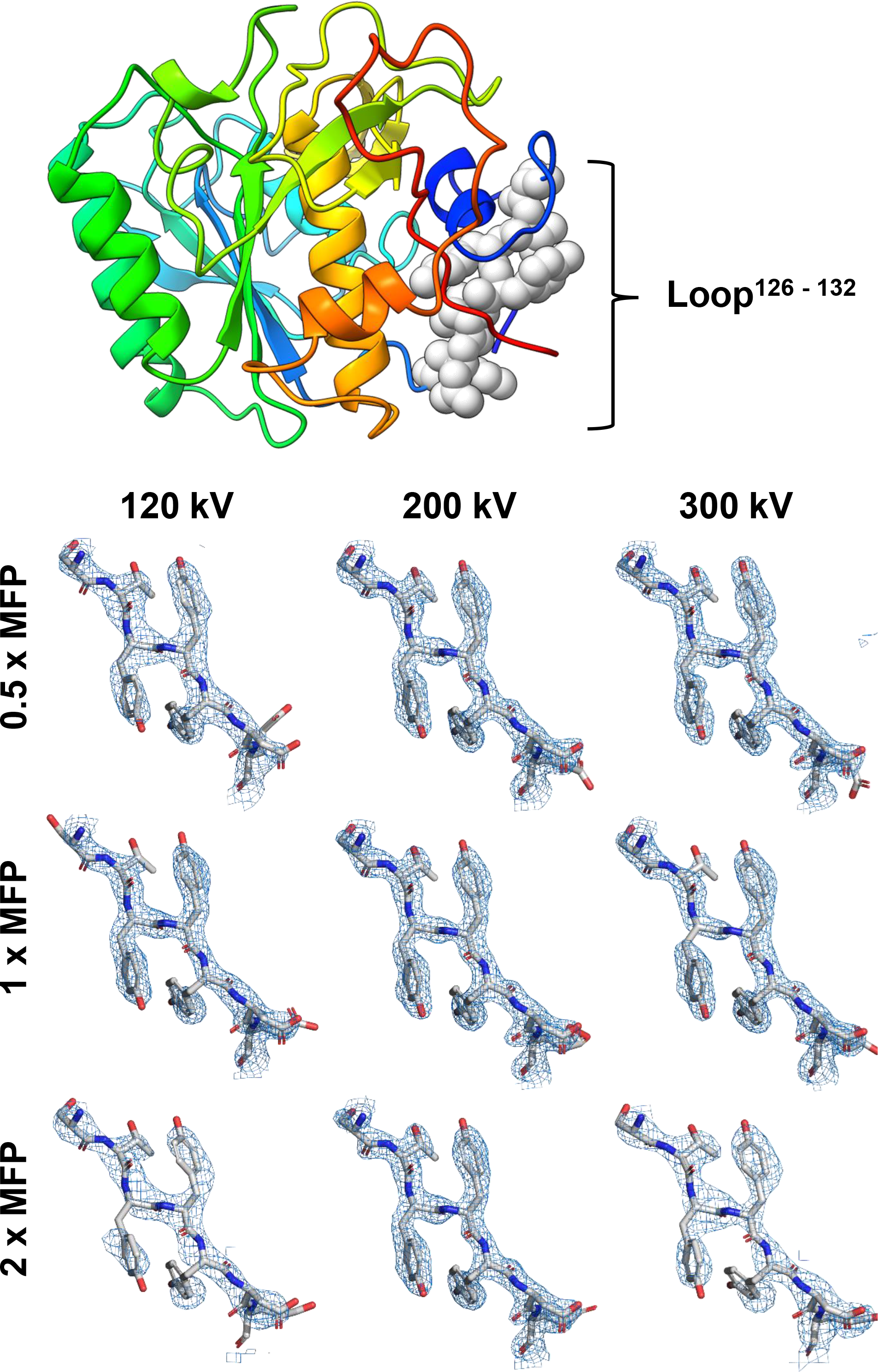
(Top) Structure of proteinase K determined by MicroED from milled lamella. (Bottom) Composite omit maps contoured at the 1.5 σ level around the loop region indicated above. Maps were generated in Phenix using electron scattering factors and all additional settings left as default.

**Supplementary Table 1.**
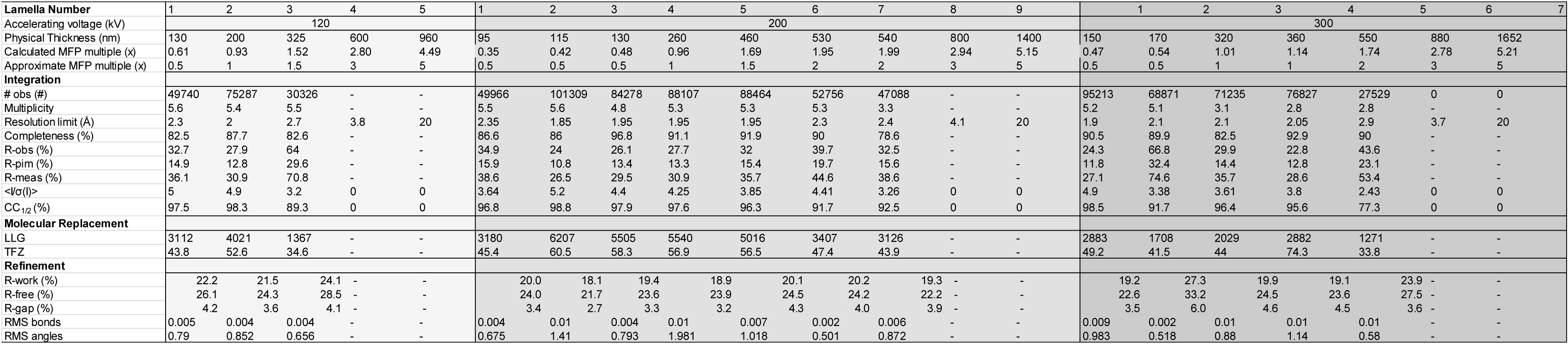
MicroED data statistics for all structures and datasets in this manuscript.

